# Disruption of the Novel Nested Gene Aff3ir Mediates Disturbed Flow-Induced Atherosclerosis in Mice

**DOI:** 10.1101/2024.10.03.616505

**Authors:** Shuo He, Lei Huang, Zhuozheng Chen, Ze Yuan, Yue Zhao, Lingfang Zeng, Yi Zhu, Jinlong He

**Affiliations:** Tianjin Key Laboratory of Metabolic Diseases, Province and Ministry Co-sponsored Collaborative Innovation Center for Medical Epigenetics, Department of Physiology and Pathophysiology, Tianjin Medical University, Tianjin, 300070, China; Department of Heart Center, The Third Central Hospital of Tianjin; Tianjin Universiy Central Hospital; Tianjin Key Laboratory of Extracorporeal Life Support for Critical Diseases; Artificial Cell Engineering Technology Research Center; Tianjin Institute of Hepatobiliary Disease; Nankai University Affinity the Third Central Hospital; Tianjin 300170, China; School of Cardiovascular and Metabolic Medicine and Sciences, King’s College London British Heart Foundation Centre of Excellence, Faculty of Life Sciences and Medicine, King’s College London, London SE5 9NU, United Kingdom

**Author notes:** **Correspondence:** Jinlong He, Ph.D. Department of Physiology and Pathophysiology, Tianjin Medical University; Qixiangtai Rd 22^nd^, Tianjin, 300070, China.; or Yi Zhu, M.D., Ph.D. Department of Physiology and Pathophysiology, Tianjin Medical University; Qixiangtai Rd 22^nd^, Tianjin, 300070, China.; or Lingfang Zeng, Ph.D. School of Cardiovascular and Metabolic Medicine and Sciences, Faculty of Life Science and Medicine, King’s College London, London SE5 9NU, United Kingdom. These authors contributed equally to this study (Shuo He, Lei Huang).

**Keywords:** Shear stress, Atherosclerosis, Nested gene, IRF5

## Abstract

Disturbed shear stress-induced endothelial atherogenic responses are pivotal in the initiation and progression of atherosclerosis, contributing to the uneven distribution of atherosclerotic lesions. This study investigates the role of Aff3ir-ORF2, a novel nested gene variant, in disturbed flow-induced endothelial cell activation and atherosclerosis. We demonstrate that disturbed shear stress significantly reduces Aff3ir-ORF2 expression in athero-prone regions. Using three distinct mouse models with manipulated AFF3ir-ORF2 expression, we demonstrate that AFF3ir-ORF2 exerts potent anti-inflammatory and anti-atherosclerotic effects in ApoE^-/-^ mice. RNA sequencing revealed that interferon regulatory factor 5 (IRF5), a key regulator of inflammatory processes, mediates inflammatory responses associated with AFF3ir-ORF2 deficiency. AFF3ir-ORF2 interacts with IRF5, promoting its retention in the cytoplasm, thereby inhibiting the IRF5-dependent inflammatory pathways. Notably, IRF5 knockdown in AFF3ir-ORF2 deficient mice almost completely rescues the aggravated atherosclerotic phenotype. Moreover, endothelial-specific AFF3ir-ORF2 supplementation using the CRISPR/Cas9 system significantly ameliorated endothelial activation and atherosclerosis. These findings elucidate a novel role for AFF3ir-ORF2 in mitigating endothelial inflammation and atherosclerosis by acting as an inhibitor of IRF5, highlighting its potential as a valuable therapeutic approach for treating atherosclerosis.

## Introduction

Atherosclerosis, characterized by the formation of fibrofatty lesions in the arterial wall, is a leading cause of morbidity and mortality worldwide, contributing to most myocardial infarctions and many strokes (Herrington, Lacey, Sherliker, Armitage, & Lewington, 2016; Libby et al., 2019). The activation of vascular endothelial cells (ECs), induced by various chemical and mechanical stimuli, such as lipopolysaccharide and shear stress, is an initial step in the development of atherosclerosis (Davignon & Ganz, 2004). Consequently, atherosclerotic lesions preferentially develop at the branches and curvatures of the arterial tree, where blood flow is disturbed (Davis, Earley, Li, & Chien, 2023). For decades, researchers have been interested in exploring the mechanisms underlying mechanotransduction during endothelial activation caused by disturbed flow (Davis et al., 2023). Several mechanosensitive proteins, such as YAP/TAZ (B. Li et al., 2019), Annexin A2 (Zhang et al., 2020), BMP4 (Sorescu et al., 2004), and NAD(P)H oxidase (Hwang et al., 2003; Jo, Song, & Mowbray, 2006), have been identified as key regulators of disturbed shear stress in ECs and have been implicated in the progression of atherosclerosis. Emerging evidence has revealed that pharmacological or genetic inhibition of endothelial YAP activation ameliorates the progression of atherosclerotic plaques in mice (B. Li et al., 2019; Wang et al., 2016; Yang et al., 2021), indicating that targeting disturbed flow-induced endothelial activation could be a promising therapeutic strategy for atherosclerosis. However, the precise mechanisms by which the disturbed flow exerts detrimental effects remain unclear.

The interferon regulatory factor (IRF) family of transcription factors, comprising nine members (IRF1-IRF9) in mammals, is primarily characterized by its role in mediating antiviral responses and type I interferon production (Sato, Taniguchi, & Tanaka, 2001). Although these members share a conserved DNA-binding domain in their N-terminal region that recognizes similar DNA sequences, IRF5 plays a central role in inflammation (Almuttaqi & Udalova, 2019; Takaoka et al., 2005). IRF5 mediates the production of proinflammatory cytokines, including IL-12b and IL-23a, and promotes the expression of inflammatory genes (Cai, Yao, & Li, 2017; Saliba et al., 2014; Weiss, Blazek, Byrne, Perocheau, & Udalova, 2013). It promotes inflammatory responses in various immune cells, including macrophages (Seneviratne et al., 2017), neutrophils (Weiss et al., 2015), and B cells (Savitsky, Yanai, Tamura, Taniguchi, & Honda, 2010). Global or myeloid-specific knockouts of IRF5 have been shown to exert anti-atherosclerotic effects (Leipner et al., 2021; Seneviratne et al., 2017). Despite the established importance of IRF5 in immune cells, its restrictively regulatory mechanism and role in shear stress-induced endothelial activation remain unknown.

We recently reported that a novel protein-coding nested gene, *Aff3ir*, contributes to endothelial maintenance by promoting the differentiation of vascular stem/progenitor cells (SPCs) into ECs (Zhao Y et al., 2024). AFF3ir-ORF2, encoded by the *Aff3ir* transcript variant 2, is predominantly expressed in the EC layer of the mouse aorta (Zhao Y et al., 2024). Notably, our recent study indicated that the overexpression of AFF3ir-ORF2 could enhance laminar flow-induced mRNA levels of essential EC markers in SPCs (Zhao Y et al., 2024), suggesting that AFF3ir-ORF2 may be a novel mechanotransduction protein in ECs. However, the regulation of Aff3ir-ORF2 under disturbed flow and its role in atherosclerosis remain unclear.

In this study, we aimed to elucidate the mechanism by which disturbed blood flow induces endothelial activation and atherosclerosis. Our study showed that disrupted Aff3ir-ORF2 expression in athero-prone regions led to inflammatory responses and development of atherosclerosis. Aff3ir-ORF2, the expression of which is reduced by disturbed shear stress, exerts critical anti-inflammatory effects by binding to IRF5 and mitigating disturbed shear stress-induced IRF5 activation. Additionally, we demonstrated that endothelial-specific supplementation with Aff3ir-ORF2 significantly ameliorated disturbed flow-induced endothelial activation and the development of atherosclerotic plaques, highlighting its promising therapeutic potential for the treatment of atherosclerosis.

## Results

### Disturbed shear stress reduces the expression of AFF3ir-ORF2

Our recent study showed the active participation of *Aff3ir* in EC differentiation from vascular SPCs induced by laminar shear stress (Zhao Y et al., 2024), suggesting the potential involvement of this novel protein-encoding nested gene in mediating hemodynamic stimulation. To further elucidate the functional role of *Aff3ir* and its encoded proteins in disturbed shear stress-induced EC activation, we examined the expression of *AFF3* and *AFF3ir* in the intima of mouse aorta. We found that the mRNA level of *AFF3ir*, but not its parent gene *AFF3*, was significantly lower in the intima of the aortic arch, an area exposed to disturbed shear stress, compared to the intima of the thoracic aorta, which was exposed to steady unidirectional shear stress (B. Li et al., 2019) (Figure 1A). *Aff3ir* transcript variants can generate two proteins (Zhao Y et al., 2024), therefore, we measured the protein levels of AFF3ir-ORF1 and AFF3ir-ORF2. While AFF3ir-ORF1 and AFF3 showed comparable expression levels in the intima of aortic arch and thoracic aorta of mice, the expression of AFF3ir-ORF2 showed an 87% reduction in the intima of aortic arch compared to the intima of thoracic aorta (Figure 1B, C), suggesting that AFF3ir-ORF2 may be a novel mechanosensitive protein that responds to disturbed shear stress. Enface immunofluorescence staining confirmed a marked reduction in AFF3ir-ORF2 expression in the inner curvature of aortic arch compared to both the outer curvature of aortic arch and the thoracic aorta (Figure 1D, E). Moreover, to demonstrate the change in AFF3ir-ORF2 within the same visual field, we examined its expression in longitudinal sections of the mouse aorta (B. Li et al., 2019). We found that the expression of AFF3ir-ORF2, but not AFF3, was notably downregulated in athero-prone regions (the inner curvature of the aortic arch and bifurcation of the carotid artery) compared to that in the protective region in the outer curvature of the aortic arch (Figure 1F). Additionally, we found that the expression of AFF3, AFF3ir-ORF1, and AFF3ir-ORF2 in the media and adventitia was comparable between the aortic arch and the thoracic aorta (Figure S1A, B).

**Figure 1.**
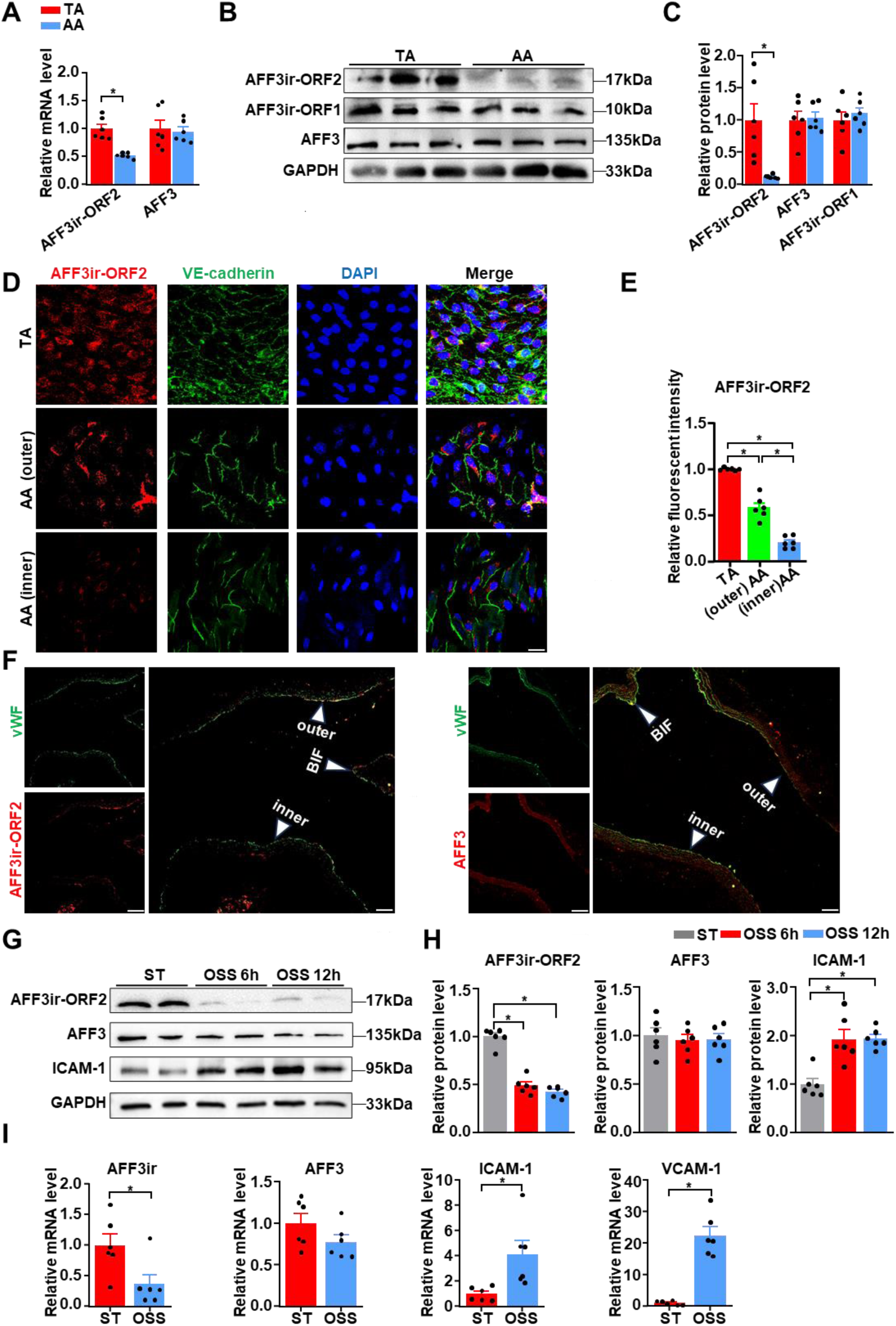
Disturbed shear stress reduces the expression of AFF3ir-ORF2 *in vivo* and *in vitro*. **A,** RT-PCR analysis of the mRNA levels of *AFF3ir-ORF2* and AF4/FMR2 family member 3 (*AFF3*) in the intima of thoracic aorta (TA) and aortic arch (AA) of C57BL/6 mice. Data are presented as mean ± SEM (n=6 mice per group). \**P*<0.05, unpaired two-tailed *t*-test. **B, C,** Western blot analysis of the expression of the indicated proteins in the intima of TA and AA of C57BL/6 mice. Protein levels were normalized to those of GAPDH, and the relative expression values were compared to those of the TA group. Data are presented as mean ± SEM (n=6 mice per group). \**P*<0.05, unpaired two-tailed *t*-test. **D, E,** En-face immunofluorescence staining of AFF3ir-ORF2, VE-cadherin, and DAPI, and quantification of AFF3ir-ORF2 expression in inner curvature of the AA (AA inner), outer curvature of the AA (AA outer), and TA of C57BL/6 mice. Scale bar, 20 μm. The immunofluorescence intensity of AFF3ir-ORF2 was normalized to that of DAPI, and the relative expression values were compared to that of the TA group. Data are presented as mean ± SEM (n=6 mice per group). \**P*< 0.05, one-way ANOVA with Tukey post-test. **F,** Representative immunofluorescent staining for von Willebrand factor (vWF), AFF3ir-ORF2, and AFF3 in longitudinal aortic sections of C57BL/6 mice. n=6 mice per group. Scale bar, 25 μm. Inner, inner curvature of the AA; outer, outer curvature of the AA; BIF, Bifurcation. **G, H,** Mouse embryonic fibroblasts (MEFs) isolated from the embryo of C57BL/6 mice were subjected to static (ST) or oscillatory shear stress (OSS, 0.5 ± 4 dyn/cm^2^, 1 Hz) for indicated time. Western blot analysis of the indicated proteins. Protein levels were normalized to GAPDH and the relative expression values were compared to that of the ST group. Data are mean ± SEM (n = 6 independent experiments). \**P*< 0.05, one-way ANOVA with Tukey post-test. **I,** MEFs were subjected to ST or OSS treatment for 6 h. RT-PCR analysis of the mRNA levels of *AFF3ir*, *AFF3,* intercellular adhesion molecule 1 (*ICAM-*1), and vascular cell adhesion molecule 1 (*VCAM-1*) in MEFs. Data are mean ± SEM (n = 6 independent experiments). \**P*< 0.05, unpaired two-tailed *t*-test.

Next, we explored the impact of disturbed shear stress on AFF3ir-ORF2 expression *in vitro*. Mouse embryonic fibroblasts (MEFs) exhibit responses consistent with those of ECs (Chen et al., 2010; Wen et al., 2013), therefore, we investigated AFF3ir-ORF2 expression in MEFs from WT mice exposed to static or disturbed flow (0.5 ± 4 dyn/cm^2^, 1 Hz). Consistent with our *in vivo* findings, while disturbed shear stress increased the expression of VCAM-1, a critical inflammatory marker of ECs (Nakashima, Raines, Plump, Breslow, & Ross, 1998), it significantly reduced both the protein and mRNA levels of AFF3ir-ORF2 (Figure 1G–I). The expression of AFF3 in response to the disturbed flow was minimally affected at both the mRNA and protein levels (Figure 1G–I). These results collectively demonstrate that disturbed shear stress induces a reduction in AFF3ir-ORF2 expression both *in vivo* and *in vitro*.

### AFF3ir-ORF2 ameliorates disturbed shear stress-induced inflammation and atherosclerosis

Disturbed shear stress-induced atherogenic responses are initial events in atherosclerotic plaque formation (Davis et al., 2023). To elucidate the regulatory role of AFF3ir-ORF2 in disturbed shear stress-induced inflammation, we overexpressed AFF3ir-ORF2 in MEFs. AFF3ir-ORF2 overexpression attenuated ICAM-1 expression induced by disturbed shear stress at both the protein and mRNA levels (Figure 2A-B, Figure S2A). To further validate our findings in ECs, we overexpressed AFF3ir-ORF2 in human umbilical vein endothelial cells (HUVECs). Consistent with our previous results, AFF3ir-ORF2 overexpression reduced the protein level of ICAM-1 induced by disturbed shear stress in HUVECs (Figure 2C-D). Moreover, AFF3ir-ORF2 overexpression attenuated disturbed shear stress-induced expression of several inflammatory genes, including VCAM-1, IL-6, and IL-1β, in both MEFs and ECs. (Figure S2A-B). Interestingly, we found that AFF3ir-ORF2 overexpression did not affect the basal expression of these inflammatory genes under ST conditions (Figure S2A-B), likely due to the relatively low levels of inflammatory gene expression under ST compared to OSS conditions. Furthermore, we measured the concentrations of inflammatory factors, including IL-6 and IL- 1β, in the culture medium of MEFs. As expected, while AFF3ir-ORF2 overexpression had little effect on the concentrations of IL-6 and IL-1β under ST condition, it significantly reduced their release induced by disturbed shear stress (Figure 2E).

**Figure 2.**
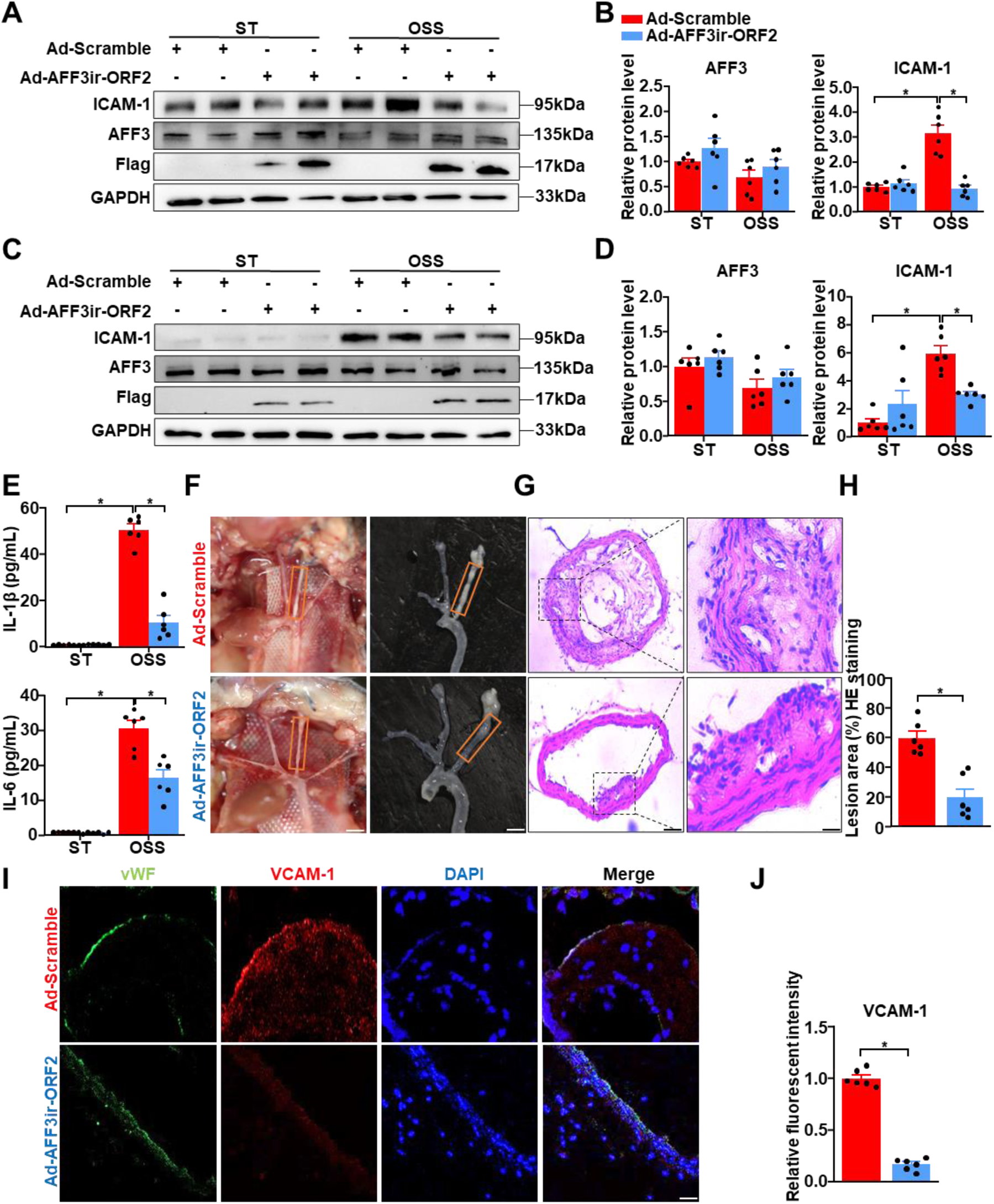
AFF3ir-ORF2 overexpression alleviates disturbed flow-induced inflammation and atherosclerosis. **A-B,** Mouse embryonic fibroblasts (MEFs) isolated from C57BL/6 mice were infected with indicated adenoviruses (Ad-Scramble or Ad-Aff3ir-ORF2) for 48 h and then exposed to static (ST) or oscillatory shear stress (OSS, 0.5 ± 4 dyn/cm^2^, 1 Hz) for another 6 h. Western blot analysis of the indicated proteins and quantification of their relative expression levels are shown. The protein levels were normalized to GAPDH and the relative expression values were compared to MEFs infected with Ad-Scramble and treated with ST. Data are presented as mean ± SEM (n = 6 independent experiments). \**P*<0.05, two-way ANOVA with Tukey post-test. **C- D,** Human umbilical vein endothelial cells (HUVECs) were infected with Ad-Scramble or Ad- AFF3ir-ORF2 for 48 h and then exposed to ST or OSS for an additional 6 h. Western blot analysis of the indicated proteins and quantification of their relative expression levels are shown. The protein levels were normalized to GAPDH and the relative expression values were compared to HUVECs infected with Ad-Scramble and treated with ST. Data are presented as mean ± SEM (n = 6 independent experiments). \**P*<0.05, two-way ANOVA with Tukey post-test. **E,** MEFs were infected with Ad-Scramble or Ad-Aff3ir-ORF2 for 48 h and then exposed to ST or OSS for another 6 h. The concentration of IL-6 and IL-1β in cell culture medium were dectected with ELISA. The relative cytokines levels is relative to MEFs infected with Ad- Scramble and treated with ST. Data are presented as mean ± SEM (n = 6 independent experiments). \**P*< 0.05, two-way ANOVA with Tukey post-test. **F-J,** Eight-week-old male ApoE^−/−^ mice were subjected to partial ligation of the carotid artery along with 10 μL of adenovirus suspension at 1 × 10^8^ transducing units (TU)/mL was instilled into the left carotid artery (LCA). The mice were then fed high-fat diet for 4 weeks. **F,** Arterial tissues were isolated to examine the atherosclerotic lesions. Scale bar, 2 mm. **G, H,** LCAs were sectioned for hematoxylin and eosin staining. Quantification of the lesion area in LCAs was shown. Scale bar, 25 μm. Data are presented as mean ± SEM (n=6 mice per group). \**P*<0.05, unpaired two-tailed *t*-test. **I**, **J,** Immunofluorescence staining for vWF, VCAM-1, and DAPI in the LCAs, and quantification of the relative fluorescent intensity of VCAM-1. The immunofluorescence intensity of VCAM-1 was normalized to DAPI, and the relative expression values were compared to that of the Ad-Scramble group. Scale bar, 50 μm. Data are presented as mean ± SEM (n=6 mice per group). \**P*<0.05, unpaired two-tailed *t*-test.

Given the anti-inflammatory effects of AFF3ir-ORF2, we speculated that it may ameliorate disturbed shear stress-induced inflammation and atherosclerosis *in vivo*. ApoE knockout (ApoE^-/-^) mice were subjected to partial ligation surgery to induce disturbed flow in the left carotid arteries (LCAs). The endothelium of the LCAs was intravascularly infected with adenovirus (Ad-Scramble or Ad-AFF3ir-ORF2) prior to surgery (Nam et al., 2009; Zhang et al., 2020). Enface immunofluorescence staining confirmed the successful AFF3ir-ORF2 overexpression in the left carotid artery (Figure S2C-D). Mice infected with Ad-AFF3ir-ORF2 exhibited a significant decrease in lesion area in the LCAs compared to those infected with Ad-Scramble (23±17% vs 63±14%) (Figure 2F-H), with no obvious plaque formation observed in the right carotid arteries (Figure S2E). Overexpression of AFF3ir-ORF2 also attenuated disturbed flow-induced inflammatory responses, as evidenced by decreased VCAM-1 expression in the endothelium of LCAs (Figure 2I-J). These findings suggested that AFF3ir-ORF2 ameliorates shear stress-induced inflammation and atherosclerosis.

### AFF3ir-ORF2 deficiency aggravates inflammation and atherosclerotic lesions in ApoE^-/-^ mice

To explore the effects of AFF3ir-ORF2 on inflammation and atherosclerosis, we generated AFF3ir-ORF2 global knockout (AFF3ir-ORF2^-/-^) mice. Genotyping PCR (Figure S3A) and western blot analysis of AFF3ir-ORF2 expression in mouse aortas (Figure S3B-C) confirmed the successful knockout. No obvious phenotypic abnormalities were observed in AFF3ir-ORF2^-/-^ mice up to 20 weeks of age and monitoring was discontinued thereafter. Additionally, AFF3ir-ORF2 deficiency did not alter systolic blood pressure, diastolic blood pressure, or mean arterial pressure (Figure S3D), suggesting that AFF3ir-ORF2 is dispensable for physiological blood pressure maintenance. We then isolated MEFs from WT and AFF3ir-ORF2^-/-^ mice. RT-PCR analysis confirmed the deficiency of AFF3ir-ORF2 in AFF3ir-ORF2^-/-^ MEFs (Figure S3E). Interestingly, the AFF3ir-ORF2 knockdown efficiency showed discrepancies between the western blot (Figure S3B) and RT-PCR results (Figure S3E). In addition to the technical differences between PCR and western blot, the characteristics of AFF3ir-ORF2 may also contribute to this inconsistency. The parent gene, AFF3, is located in a genetically variable region, and it can be excised via intron 5 to form a replicable transposon that translocates to other chromosomes, potentially contributing to leukemia (Chinen et al., 2008; Hiwatari et al., 2003; Miller, Leventaki, Harker-Murray, Drendel, & Bone, 2022; von Bergh et al., 2002). AFF3ir, located in intron 6, exists within this transposon, which may complicate the measurement of its expression. Furthermore, we found that AFF3ir-ORF2 deficient MEFs displayed higher expression of inflammatory genes, including *ICAM-1, VCAM-1,* and *IL-1b*, compared to those in WT MEFs, under disturbed flow stimulation (Figure S3F).

Next, we crossed AFF3ir-ORF2^-/-^ mice with ApoE^-/-^ mice to generate double-knockout (ApoE^-/-^AFF3ir-ORF2^-/-^) mice. Eight-week-old ApoE^-/-^ and ApoE^-/-^AFF3ir-ORF2^-/-^ mice were fed a high-fat diet for 12 weeks to induce atherosclerosis (Figure 3A). *En face* Oil-Red O staining indicated that AFF3ir-ORF2 deficiency accelerated the development of atherosclerosis in the entire aorta, AA, and TA. (Figure 3B, C). Furthermore, AFF3ir-ORF2 deletion increased the lesion area and lipid deposition in the aortic roots of ApoE^-/-^ mice without altering the collagen fiber content (Figure 3D, E). Similar results were observed in distributing arteries (LCAs) (Figure 3F, G). Given that the expression of adhesion proteins, such as VCAM-1 in ECs is crucial for monocyte infiltration into plaques (Kobiyama & Ley, 2018), we assessed VCAM-1 expression in the aortic roots of these mice. We found that AFF3ir-ORF2 deletion increased VCAM-1 expression in the aortic roots of ApoE^-/-^ mice (Figure 3H, I), indicating that the atherogenic effects of AFF3ir-ORF2 deletion may result from endothelial inflammation. Additionally, there were no significant differences between the two groups in body weight or triglyceride, total cholesterol, LDL cholesterol, and HDL cholesterol levels (Figure S4A, B), indicating that the atherogenic effect of AFF3ir-ORF2 silencing is unlikely to be related to lipid metabolism. Taken together, these results indicate that AFF3ir-ORF2 deficiency aggravates inflammation and atherosclerotic lesions in mice.

**Figure 3.**
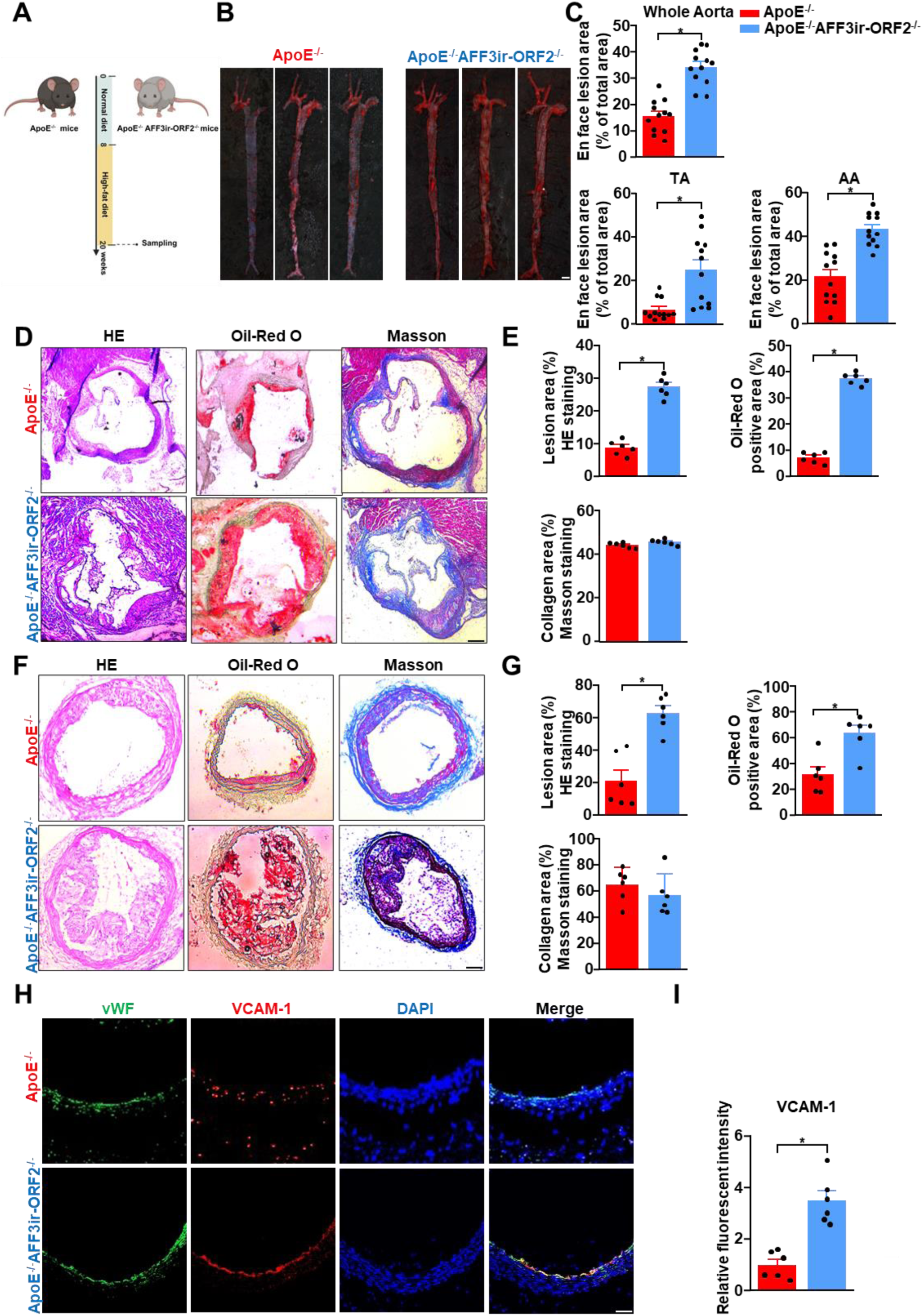
AFF3ir-ORF2 deletion aggravates inflammation and atherosclerotic lesions in ApoE^-/-^ mice. Eight-week-old male ApoE^-/-^ and ApoE^-/-^AFF3ir-ORF2^-/-^ mice were fed a high-fat diet for 12 weeks. Arterial tissues and aortic roots were isolated to examine atherosclerotic lesions. **A,** Schematic of experimental strategy. **B,** Representative images of en face Oil-Red O staining of the aortas. Scale bar, 4 mm. **C,** Quantification of the plaque area in the whole aorta, aortic arch (AA), and thoracic aorta (TA). Data are presented as mean ± SEM (n=12 mice per group). \**P*<0.05, unpaired two-tailed *t*-test. **D,** Oil-Red O, hematoxylin and eosin (HE), and Masson staining of the aortic roots. Scale bars, 500 μm. **E,** Quantification of plaque size, Oil-Red O- positive area, and collagen fiber content in aortic root sections. Data are presented as mean ± SEM (n=6 mice per group). \**P*< 0.05, unpaired two-tailed *t*-test. **F**, LCAs were sectioned and stained with Oil-Red O, HE, and Masson’s trichrome. Scale bars, 500 μm. **G,** Quantification of plaque size, Oil-Red O-positive area, and collagen fiber content in the LCA sections. Data are presented as mean ± SEM (n=6 mice per group). \**P*<0.05, unpaired two-tailed *t*-test. **H,** Representative immunofluorescence images of vWF, VCAM-1, and DAPI in the aortic roots. Scale bar, 500 μm. **I**, Quantification of the relative fluorescence intensity of VCAM-1. The immunofluorescence intensity of VCAM-1 was normalized to that of DAPI, and the relative expression values were compared to that of the ApoE^-/-^ group. Data are presented as mean ± SEM (n=6 mice per group). \**P*<0.05, unpaired two-tailed *t*-test.

### AFF3ir-ORF2 mitigates disturbed shear stress-induced inflammation by interacting with IRF5 and retaining it within the cytosol

To explore the mechanism by which AFF3ir-ORF2 mitigates atherogenesis, we performed RNA sequencing (RNA-seq) on MEFs from WT and AFF3ir-ORF2^-/-^ mice. The Principal component analysis plot depicted clear clustering of WT versus AFF3ir-ORF2^-/-^ samples (Figure 4A). We identified 1,167 upregulated and 310 downregulated genes in the AFF3ir-ORF2^-/-^ group, with a criterion of 1.5-fold change and P<0.05 (Figure 4B, Figure S5A). All the differentially expressed genes were subjected to bioinformatics enrichment analysis using Gene Ontology (GO) databases. GO analysis showed that these genes were mainly enriched in processes, including leukocyte cell-cell adhesion, regulation of cell−cell adhesion, and leukocyte activation involved in immune response (Figure 4C), which is highly consistent with the phenotypes observed in AFF3ir-ORF2^-/-^ mice. To further investigate the function features of these differentially expressed genes in the context of the atherosclerotic microenvironment, we mapped the differential gene list onto the atherosclerosis-related gene dataset (Rouillard et al., 2016), resulting in 363 overlapping genes. GO analysis of these genes revealed enrichment in processes related to cell−cell adhesion and leukocyte activation involved in immune response (Figure S5B), which is highly consistent with the observed effects of AFF3ir-ORF2 on VCAM- 1 expression.

**Figure 4.**
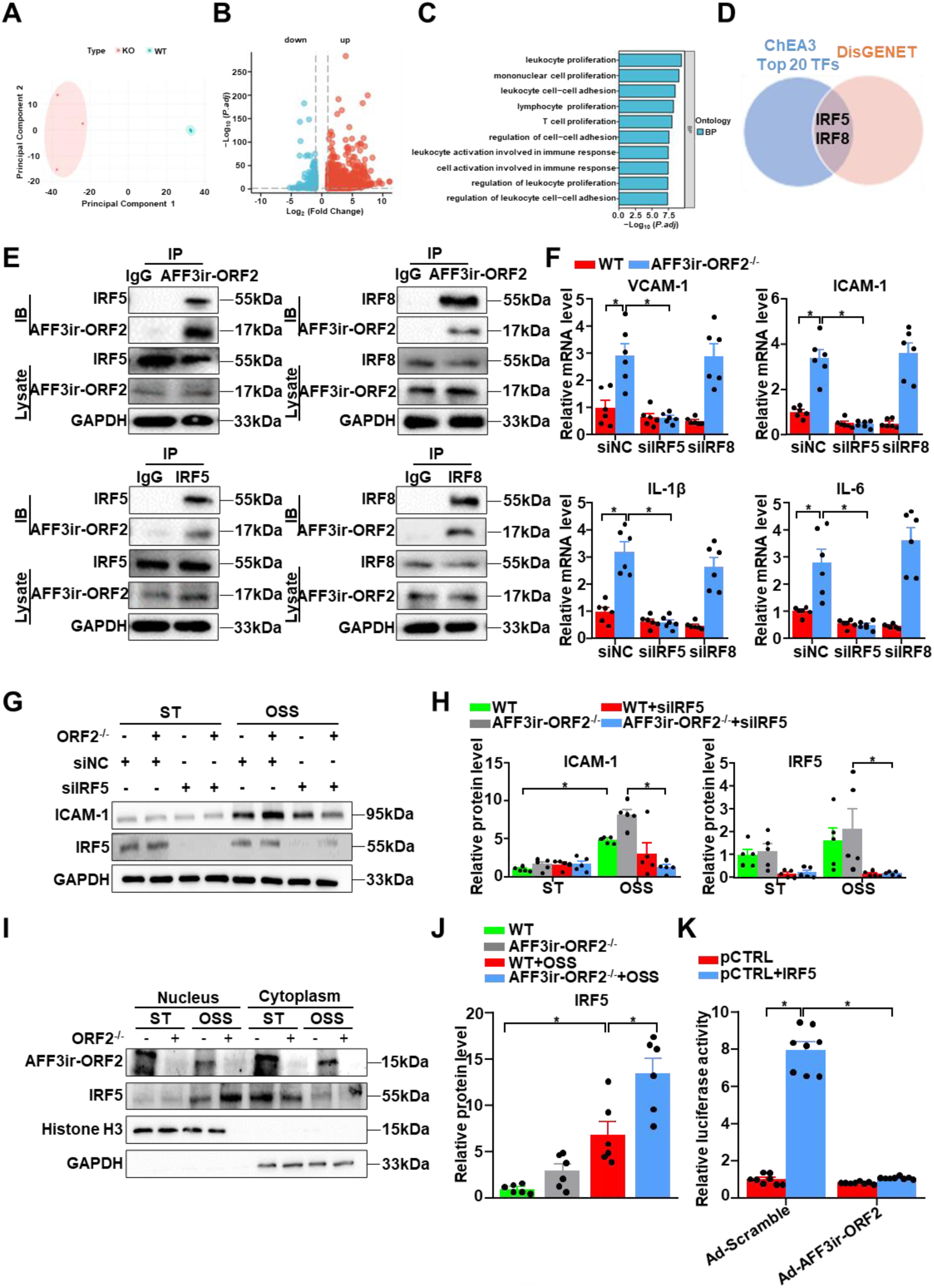
AFF3ir-ORF2 mitigates disturbed shear stress-induced inflammation by interacting with IRF5 and retaining it within the cytosol. **A–D,** Mouse embryonic fibroblasts (MEFs) were isolated from wild-type (WT) and AFF3ir-ORF2^-/-^ mice. **A,** Principal component analysis (PCA) analysis of RNA-seq data to visualize sample to sample variation. **B**, Volcano map showing mRNA profiles of WT and AFF3ir-ORF2^-/-^ MEFs (n=3). **C,** Gene Ontology enrichment pathway analysis of the differentially expressed genes. **D**, Venn diagrams of the top 20 transcription factors from the ChEA3 and DisGENET analysis related to atherosclerosis. **E,** Immunoprecipitation performed using antibodies against AFF3ir-ORF2, IRF5, and IRF8. n = 3 independent experiments. **F–H,** WT and AFF3ir-ORF2^-/-^ MEFs were subjected to silence of Control (siNC), IRF5 (siIRF5), or IRF8 (siIRF8) with siRNAs for 24 h, followed by exposure to static (ST) or oscillatory shear stress (OSS, 0.5 ± 4 dyn/cm^2^, 1 Hz) for another 6 h. **F**, RT-PCR analysis of the mRNA levels of VCAM-1, ICAM-1, IL-6, and IL-1β. The relative expression values were compared to WT MEFs transfected with siNC and treated with ST. Data are mean ± SEM (n = 6 independent experiments). \**P*< 0.05, two-way ANOVA with Tukey post-test. **G, H**, Representative western blots of IRF5 and ICAM-1 expression. Data are mean ±SEM (n = 5 independent experiments). \**P*< 0.05, two-way ANOVA with Tukey post-test. **I, J**, WT, and AFF3ir-ORF2^-/-^ MEFs were exposed to ST or OSS for 6 h. Nuclear and cytoplasmic proteins were extracted from the cells. Representative western blots of the indicated proteins and quantification of IRF5 expression in nucleus are shown. The expression of these proteins was relative to the level of nuclear IRF5 in ST-treated WT MEFs. Data are mean ± SEM (n = 6 independent experiments). \**P*< 0.05, one-way ANOVA with Tukey post-test. **K,** HEK293 cells were transfected with the firefly luciferase reporter plasmid containing the IRF5-responsive ZNF217 promoter along with a β- galactosidase reporter plasmid for 24 hours. Cells were infected with the indicated adenoviruses (Ad-Scramble or Ad-Aff3ir-ORF2) for 24 h. Promoter activity was measured using luciferase, which was normalized to β-gal. Data are mean ±SEM (n = 6 independent experiments). \**P*< 0.05, two-way ANOVA with Tukey post-test.

To further identify the upstream transcriptional regulators of these genes, we used the list of differentially expressed genes from the RNA-seq data to predict upstream transcription factors using the ChEA3 database (Keenan et al., 2019). Then, the top 20 transcription factors obtained from the ChEA3 database were mapped to the atherosclerotic disease-related gene list in the Disgenet database (Pinero et al., 2017). Interferon regulatory factor 5 (IRF5) and IRF8 were identified as key upstream regulators (Figure 4D). IRF5 and IRF8, which are members of the same family of transcription factors originally implicated in interferon production, have been identified as critical regulators of the inflammatory response and contribute to the pathogenesis of various inflammatory diseases (Almuttaqi & Udalova, 2019; Salem, Salem, & Gros, 2020). However, their potential roles in disturbed shear stress-induced inflammation remain unclear. We speculated that AFF3ir-ORF2 interacts with IRF5 and/or IRF8. Coimmunoprecipitation assays indicated that endogenous AFF3ir-ORF2 could bind to both IRF5 and IRF8 (Figure 4E). To determine which transcription factor that mediates the inflammatory effects of AFF3ir-ORF2 deficiency, we silenced IRF5 and IRF8 in WT and AFF3ir-ORF2^-/-^ MEFs exposed to disturbed flow. Notably, silencing IRF5, but not IRF8, blunted the upregulation of inflammatory genes, including *ICAM-1, VCAM-1, IL-6,* and *IL-1β* (Figure 4F), suggesting that IRF5 was the predominant factor mediating the anti-inflammatory effects of AFF3ir-ORF2 in the context of disturbed shear stress. Consistently, we found that IRF5 silencing significantly inhibited the upregulation of ICAM-1 protein levels induced by AFF3ir-ORF2 deficiency under disturbed shear stress (Figure 4G, H). In addition, neither IRF5 nor IRF8 expression levels were affected by AFF3ir-ORF2 deficiency (Figure S5C). However, we found that AFF3ir-ORF2 deficiency significantly increased the expression of IRF5-targeted genes (predicted by the ChEA3 database), including ICAM-1, CCL5, and CXCL10 (Figure S5D). Notably, the protein level of IRF5 was not significantly affected by disturbed shear stress (Figure 4G, H). Consistently, the mRNA levels of IRF5 were previously reported to be barely changed in the context of disturbed shear stress (GSE276195, Figure S5E) or the atherosclerotic environment (GSE222583, Figure S5F). Given that the transcriptional activity of IRF5 depends on its nuclear translocation (Lv et al., 2024), we next explored whether AFF3ir-ORF2 affects the subcellular localization of IRF5. Subcellular fractionation assays indicated that IRF5 was predominantly localized in the cytoplasm under static conditions, but exhibited obvious nuclear localization when exposed to disturbed shear stress (Figure 4I, J). While the total expression of IRF5 was barely affected by AFF3ir-ORF2 deficiency or overexpression, nuclear localization of IRF5 increased with AFF3ir-ORF2 deficiency (Figure 4I, J). To further ascertain the role of AFF3ir-ORF2 in regulating the transcriptional activity of IRF5, we performed a luciferase reporter assay (Qiao, Lv, Qiao, Wang, & Miao, 2022). AFF3ir-ORF2 overexpression significantly decreased the transcriptional activity of IRF5 (Figure 4K). In summary, these results suggested that AFF3ir-ORF2 acts as an endogenous inhibitor of IRF5 and exerts anti-inflammatory effects by retaining IRF5 in the cytosol.

### IRF5 knockdown prevents the aggravation of atherosclerosis induced by ORF2 deficiency

Next, we investigated the role of IRF5 in disturbed flow-induced atherosclerosis *in vivo* and whether it mediates the atherogenic phenotype associated with AFF3ir-ORF2 deficiency. ApoE^-/-^ and ApoE^-/-^AFF3ir-ORF2^-/-^ mice were subjected to partial ligation surgery in the LCAs and intravascularly infected with lentiviruses expressing either IRF5-specific shRNA (lenti-shIRF5) or Scramble shRNA (lenti-shScramble). En-face immunofluorescence staining confirmed successful IRF5 deletion in the left carotid artery (Figure S6A). After a 4-week high-fat diet challenge, IRF5 deletion resulted in an approximately 60% reduction in plaque area in the LCAs of ApoE^-/-^ mice (Figure 5A, B). In addition, IRF5 deletion attenuated endothelial activation, as evidenced by reduced VCAM-1 expression in the endothelium of LCAs (Figure 5C, D). Notably, although ApoE^-/-^AFF3ir-ORF2^-/-^ mice exhibited an increased plaque area in the LCAs compared to ApoE^-/-^ mice, IRF5 deletion almost completely abolished these differences, reducing both the plaque area and VCAM-1 expression in the endothelium of the LCAs (Figure 5A–D). These findings provide in vivo evidence that AFF3ir-ORF2 deficiency-induced atherosclerosis is mediated by endothelial IRF5.

**Figure 5.**
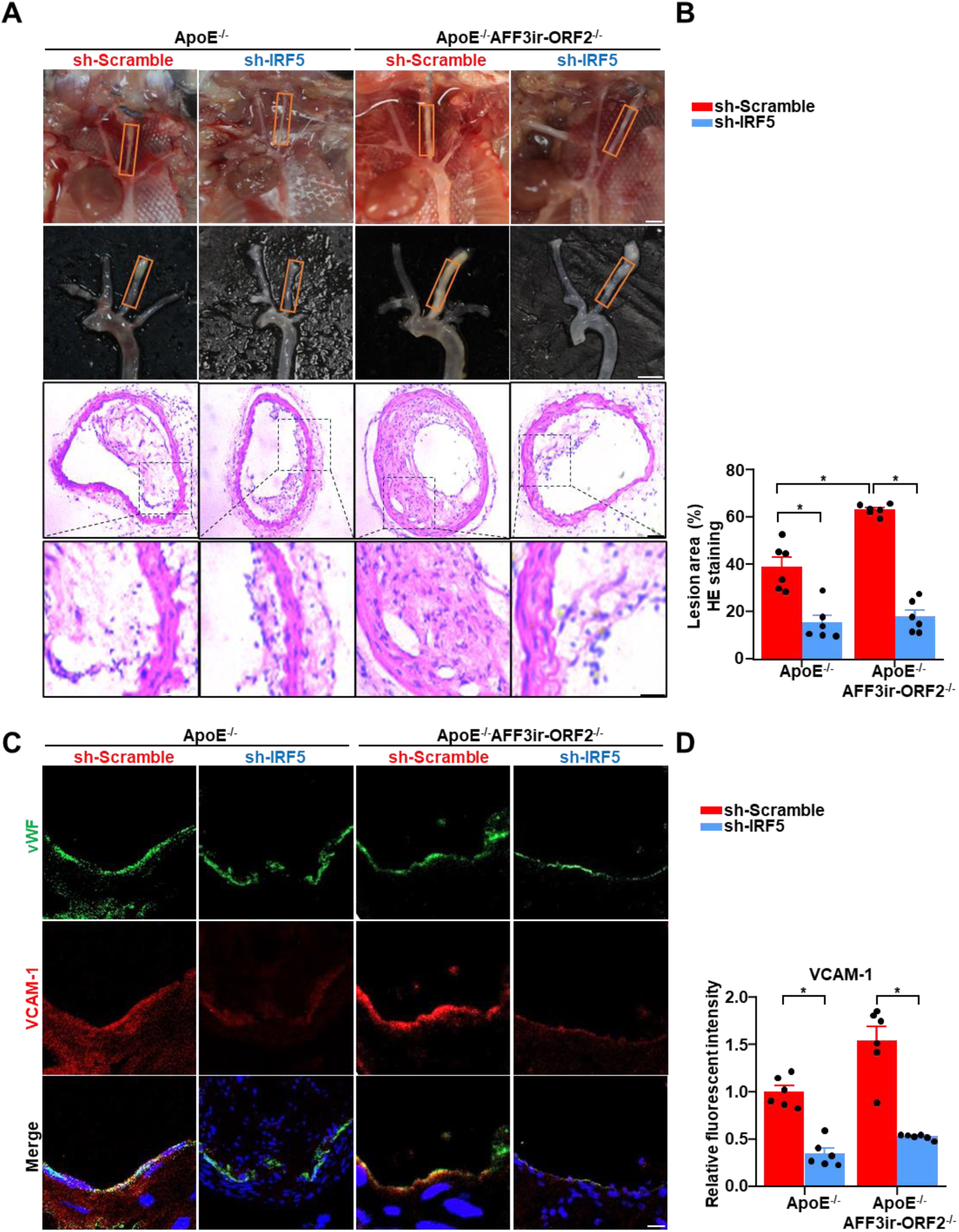
IRF5 knockdown prevents the aggravation of atherosclerosis in AFF3ir-ORF2 deficienct mice. Eight-week-old male ApoE^-/-^ mice were subjected to partial ligation of the left carotid artery (LCA) along with 10 μL of lentivirus suspension at 1 × 10^8^ transducing units (TU)/mL was instilled into the LCA. The mice were then fed a high-fat diet for 4 weeks. **A-B,** Arterial tissues were isolated to examine the atherosclerotic lesions. LCAs were sectioned for hematoxylin and eosin staining. Quantification of the lesion area in LCAs was shown. Data are mean ± SEM (n=6 mice per group). \**P*< 0.05, two-way ANOVA with Tukey post-test. Scale bar: 2 mm for gross images, 25 μm for staining images. **C, D,** Immunofluorescence staining for vWF, VCAM- 1, and DAPI in the LCAs and quantification of the relative fluorescent intensity of VCAM-1. Scale bar, 50 μm. The immunofluorescence intensity of VCAM-1 was normalized to DAPI, and the relative expression values were compared to that of the group of ApoE^-/-^ mice infected with Ad-scramble. Data are presented as mean ± SEM (n=6 mice per group). \**P*< 0.05, two-way ANOVA with Tukey post-test.

### Endothelial-specific AFF3ir-ORF2 supplementation ameliorates EC activation and atherosclerosis in mice

Given the significant anti-inflammatory effects of AFF3ir-ORF2 on endothelial activation and atherosclerosis, we explored the potential use of gene therapy targeting AFF3ir-ORF2 to treat atherosclerosis. Endothelial-specific AFF3ir-ORF2 overexpression was achieved using an EC- enhanced AAV-mediated CRISPR/Cas9 genome-editing system controlled by an EC-specific ICAM2 promoter as we previously reported (Z. Y. Li et al., 2024; Swiech et al., 2015). ApoE^-/-^ mice infected with AAV-ICAM2-Control or AAV-ICAM2- AFF3ir-ORF2 were fed a high-fat diet for 12 weeks (Figure 6A). En-face immunofluorescence staining confirmed successful AFF3ir-ORF2 overexpression in ECs (Figure S7A-B). Endothelial-specific AFF3ir-ORF2 overexpression had a minimal effect on triglycerides, total cholesterol, LDL cholesterol, and HDL cholesterol levels in the plasma of mice (Figure S7C). However, compared to the negative control, endothelial-specific AFF3ir-ORF2 overexpression significantly reduced the Oil-red O-positive lesion area in the whole aortas of ApoE^-/-^ mice (19±5% vs 54±8%) (Figure 6B, C). Moreover, endothelial-specific AFF3ir-ORF2 overexpression reduced the lesion area and lipid deposition in the aortic roots of ApoE^-/-^ mice without altering the collagen fiber content (Figure 6D, E). In addition, AFF3ir-ORF2 overexpression effectively suppressed VCAM-1 expression in the endothelium of the aortic roots of ApoE^-/-^ mice (Figure 6F, G). Collectively, these results suggest that supplementation with AFF3ir-ORF2 was effective in preventing atherosclerosis development.

**Figure 6.**
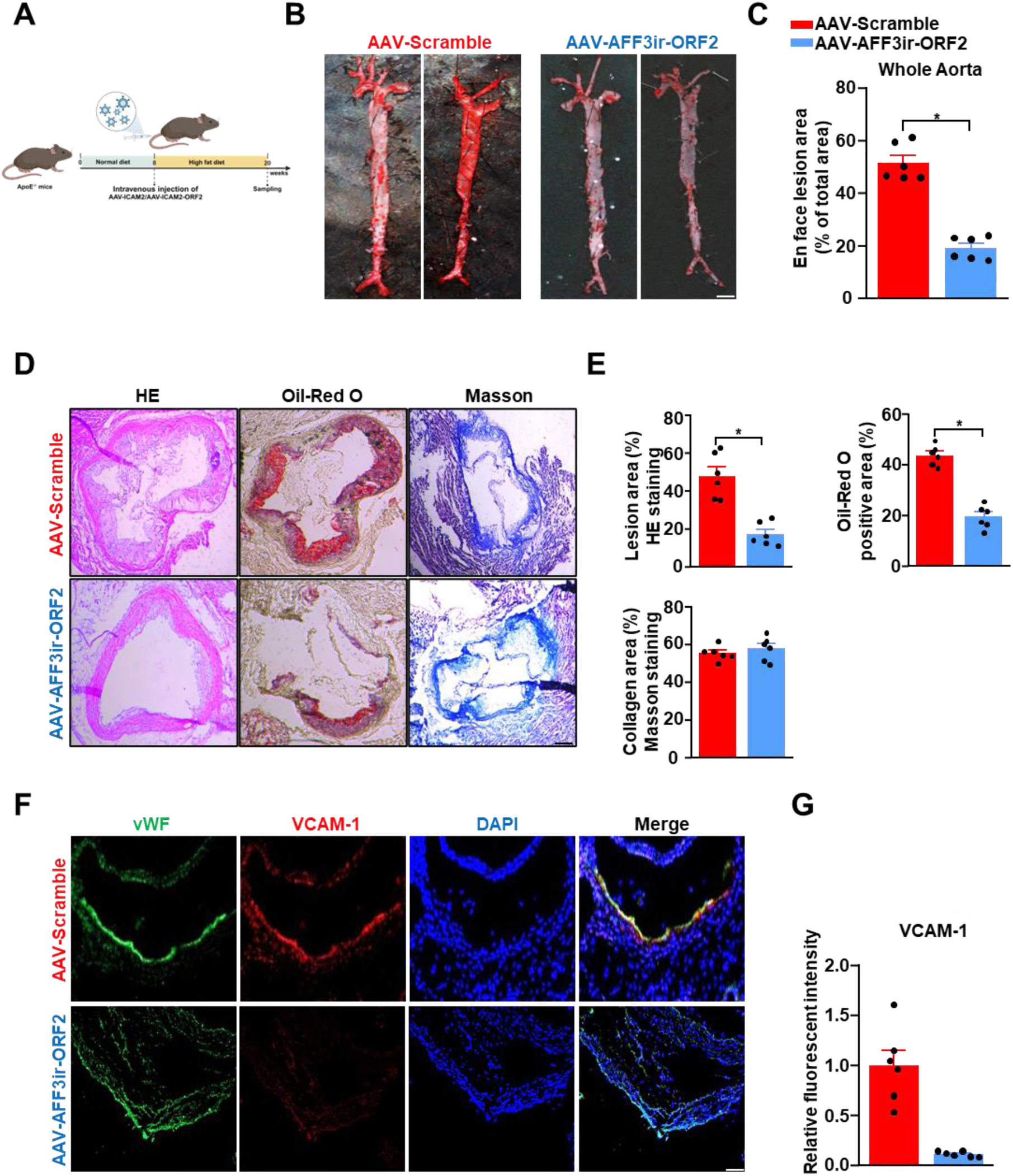
Endothelial-specific AFF3ir-ORF2 supplementation alleviates EC activation and atherosclerosis in ApoE^-/-^ mice. Eight-week old ApoE^-/-^ male mice were infused with the indicated adeno-associated virus (AAV) and then fed a high-fat diet for 12 weeks. **A**, Schematic of the experimental strategy. **B**, Representative images of enface Oil-Red O staining of the aortas. Scale bar, 4 mm. **C,** Quantification of the plaque area in the entire aortas. Data are presented as mean ± SEM (n=6 mice per group). \**P*< 0.05, unpaired two-tailed *t*-test. **D,** Hematoxylin and eosin (HE), Oil-Red O, and Masson staining of the aortic roots. Scale bars, 500 μm. **E,** Quantification of plaque size, Oil-Red O-positive area, and collagen fiber content in aortic root sections. Data are presented as mean ± SEM (n=6 mice per group). \**P*< 0.05, unpaired two-tailed *t*-test. **F**, Representative immunofluorescence image of vWF, VCAM-1, and DAPI in the aortic roots. Scale bar, 500 μm. **G**, Quantification of the relative fluorescent intensity of VCAM-1. The immunofluorescence intensity of VCAM-1 was normalized to that of DAPI, and the relative expression values were compared to that of the AAV-Scramble group. Data are presented as mean ± SEM (n=6 mice per group). \**P*< 0.05, unpaired two-tailed *t*-test.

## Discussion

Endothelial activation is a critical initial event in the development of atherosclerosis, and emerging evidence suggests that targeting disturbed shear stress-induced endothelial activation is a promising therapeutic strategy. In the present study, we elucidated the role of the novel nested gene-encoded protein, AFF3ir-ORF2, in sensing disturbed shear stress. Moreover, we demonstrated that AFF3ir-ORF2 acts as an endogenous inhibitor of IRF5, a key regulator of the inflammatory response, thereby exerting potent anti-inflammatory and anti-atherogenic effects (Figure 7).

**Figure 7.**
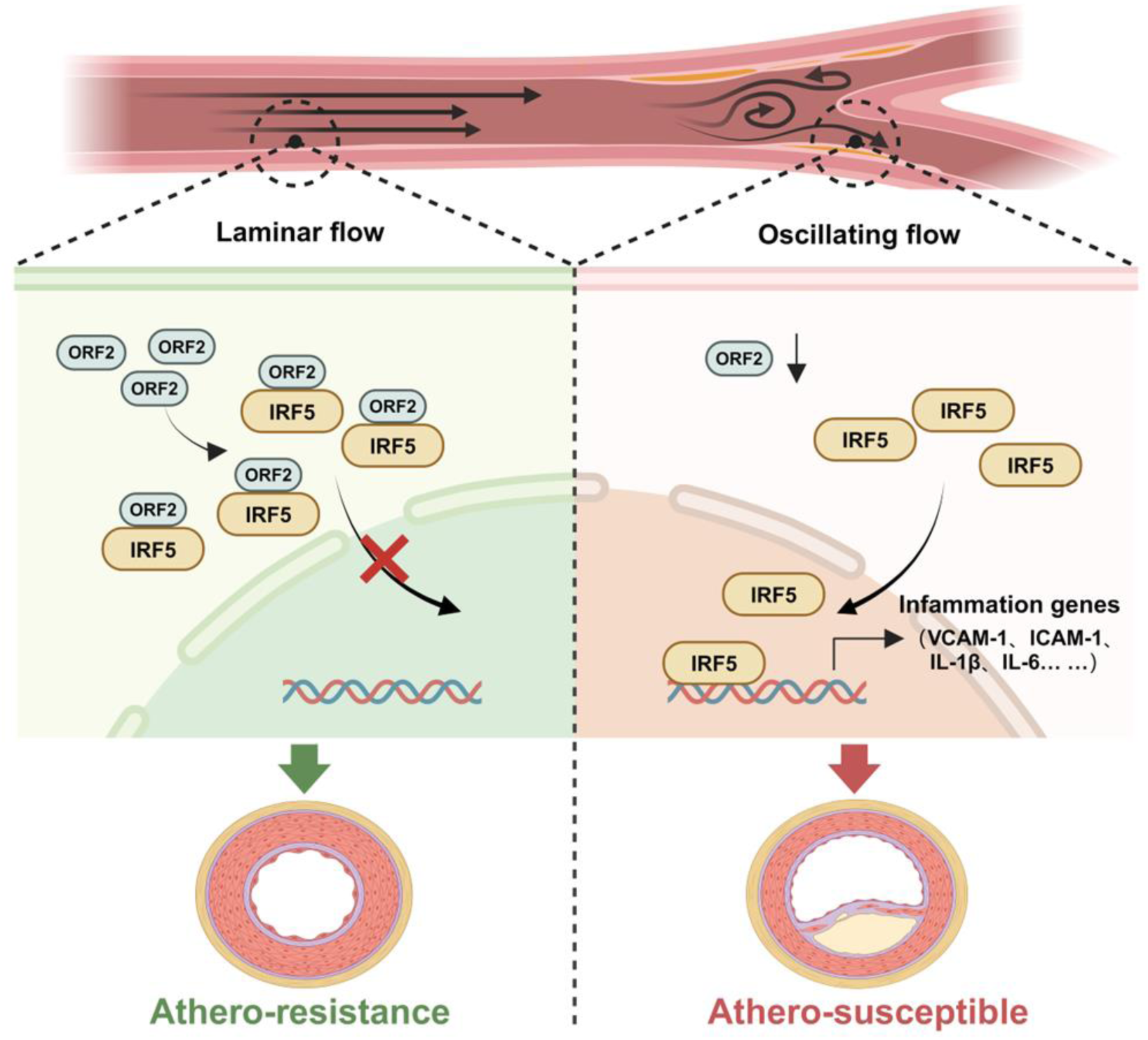
Schematic illustration of the AFF3ir-ORF2/IRF5 cascade in disturbed flow-induced endothelial activation and atherosclerosis. Disturbed flow induced a down-regulation of AFF3ir-ORF2, which could interact with IRF5 and promote the latter’s retention in the cytoplasm, thereby boosting IRF5-dependent inflammatory pathways in endothelial cells and leading to atherogenesis.

Using three mouse models (global AFF3ir-ORF2 knockout, locally AFF3ir-ORF2 endothelial expression, and endothelial-specific AFF3ir-ORF2 overexpression), we demonstrated that AFF3ir-ORF2 exerted potent anti-inflammatory and anti-atherosclerosis effects in ApoE^-/-^ mice. Notably, while AFF3ir-ORF2 knockout increased VCAM-1 expression in the endothelium and enlarged the plaque area in the aortic roots, it had a minimal effect on collagen deposition within the plaques. This discrepancy may be attributed to the differential expression patterns of IRF5 across various cell types (Roberts, Collado, & Barnes, 2024). Phenotypically modulated vascular smooth muscle cells (VSMCs) within the fibrous cap produce extracellular matrix molecules critical for plaque composition and stabilization (Bennett, Sinha, & Owens, 2016). A previous study has shown minimal colocalization between IRF5 and the VSMC marker, α- smooth muscle actin, in aortic root lesions of ApoE^-/-^ mice (Seneviratne et al., 2017), indicating a relatively low IRF5 expression level in VSMCs. Consequently, AFF3ir-ORF2 likely exerts its anti-inflammatory effects primarily through endothelial IRF5. Consistently, endothelial-specific AFF3ir-ORF2 overexpression reduced the aortic plaque area, but had minimal effects on collagen deposition. Our findings establish the potent anti-atherosclerotic role of AFF3ir-ORF2 in early and advanced atherosclerosis mouse models. However, given the multiple critical roles of ECs throughout the initiation and progression of atherosclerosis, further investigations are needed to explore the potential role of AFF3ir-ORF2 in other atherosclerotic processes, including endothelial-to-mesenchymal transition, plaque rupture, and atherothrombotic occlusion. Additionally, we found that disturbed shear stress transcriptional downregulated the expression of AFF3ir. However, the protein levels of AFF3ir-ORF2, but not those of AFF3ir-ORF1, were reduced by disturbed shear stress. Since both AFF3ir-ORF1 and AFF3ir-ORF2 are derived from AFF3ir, different translation mechanisms may be involved in the production of AFF3ir-ORF1/2.

In addition to ECs, various other cell types, particularly immune cells, play crucial roles in the progression of atherosclerotic plaques (Wolf & Ley, 2019). Although our results indicated the potent anti-inflammatory role of AFF3ir-ORF2 in ECs, the potential contributions of other cell types may also be involved, which could be further elucidated using AFF3ir-ORF2 tissue-specific knockout or overexpression mouse models. Macrophage polarization and inflammatory responses accelerate plaque development, leading to an increase in necrotic core and vulnerable plaques (Krausgruber et al., 2011). Elevated IRF5 expression and nuclear localization have been observed in macrophages within plaques of ApoE^-/-^ mice (Seneviratne et al., 2017). IRF5 has been demonstrated to drive macrophages towards a pro-inflammatory state, thereby affecting plaque stability (Seneviratne et al., 2017). Global or myeloid cell-specific deletion of IRF5 stabilizes atherosclerotic plaques by suppressing the inflammatory phenotypes of macrophages (Leipner et al., 2021; Seneviratne et al., 2017). Despite these findings, the role of IRF5 in shear stress-induced endothelial activation remains largely unknown. Our study provides evidence that disturbed shear stress is sufficient to induce IRF5 nuclear translocation and activation in ECs. Furthermore, IRF5 knockdown in ECs significantly reduced disturbed flow-induced plaque formation in LCAs. These findings suggest that targeting endothelial IRF5 may be an effective strategy for combating the early stages of atherosclerosis.

Given the key role of IRF5 in mediating inflammatory responses, it has been considered as an attractive therapeutic target, and various strategies have been developed to study and modulate its function (Almuttaqi & Udalova, 2019). For example, nanoparticle-delivered siRNA targeting IRF5 in macrophages promotes inflammation resolution, improves infarct healing, and attenuates post-myocardial infarction remodeling (Courties et al., 2014). Additionally, manipulating IRF5 protein levels through the E3 ubiquitin ligase, TRIM21, has been explored as a strategy for modulating its activity (Lazzari et al., 2014). Given the crucial physiological role of IRF5, strategies aimed at suppressing its pathophysiological activation without altering basal levels may offer additional benefits. Our study introduces a novel approach to inhibit IRF5 activation. We found that the novel nested gene-encoded protein, AFF3ir-ORF2, interacts with IRF5, leading to cytoplasmic retention and inactivation under disturbed shear stress conditions. Importantly, endothelium-specific supplementation with AFF3ir-ORF2 effectively attenuated endothelial activation and reduced the atherosclerotic plaque area in ApoE^-/-^ mice, suggesting that targeting endothelial IRF5 activation with AFF3ir-ORF2 holds promise for the treatment of atherosclerosis. Furthermore, as emerging studies have highlighted the substantial contributions of IRF5 to autoimmune diseases (Graham et al., 2006), neuropathic pain (Masuda et al., 2014), obesity (Dalmas et al., 2015), and hepatic fibrosis (Alzaid et al., 2016), future research should investigate whether AFF3ir-ORF2 has beneficial effects in these contexts.

### Conclusion

In conclusion, this study provides novel evidence that the disruption of AFF3ir-ORF2 expression under disturbed flow promotes endothelial inflammatory responses and atherosclerosis. AFF3ir-ORF2 serves as an endogenous inhibitor of IRF5 by binding to IRF5 and preventing its nuclear translocation. Supplementation with endothelial AFF3ir-ORF2 may be a promising therapeutic strategy for treating atherosclerosis.

## Materials and methods

### Animals

C57BL/6 and Apolipoprotein E-null (ApoE^-/-^) mice were purchased from the Experimental Animal Centre of Military Medical Science Academy (Beijing, China). AFF3ir-ORF2- heterozygote (AFF3ir-ORF2^+/-^) mice were acquired from Dr. Lingfang Zeng’s Laboratory at King’s College London. Briefly, gRNAs targeting the mouse AFF3ir-ORF2 locus (gRNA1: GCAACCCACGGAGTTGCAGTTGG; gRNA2: GTCATTAACTCCTTTAATATAGG; gRNA3: TGCAACTCCGTGGGTTGCTGTGG; gRNA4: GACCACACATAACAGTGAATAGG) and Cas9 mRNA were co-injected into fertilized mouse eggs to generate targeted knockout offspring. F0 founder animals were identified by PCR followed by sequence analysis, and then bred to WT mice to test germline transmission and produce F1 animals. Heterozygous targeted mice were intercrossed to generate homozygous targeted mice. The genotyping primer used were: 5’- GGAAAGACCACAGAATCAATGACA-3’, 5’-AACATTGCTATACCCCACTATA-3’. To generate ApoE and AFF3ir-ORF2 double knockout mice (ApoE^-/-^AFF3ir-ORF2^-/-^), ApoE^-/-^ mice were crossed with AFF3ir-ORF2^-/-^ mice. The animals were maintained at 21 ± 1°C under a 12-hour light/dark cycle (lights on at 07:00, lights off at 19:00) with ad libitum access to water and standard chow unless specified otherwise. This study adhered to the *Guide for the Care and Use of Laboratory Animals* of the US National Institutes of Health (NIH Publication No. 85- 23, revised 2011). All study protocols were approved by the Institutional Animal Care and Use Committee of Tianjin Medical University.

### Carotid artery partial ligation surgery

The surgery was performed as we previously described (Yang et al., 2021). Briefly, mice were anesthetized with isoflurane (2-3%). A ventral midline incision (4-5 mm) was made in the neck and the left carotid artery was exposed through blunt dissection of subcutaneous fat and muscle tissue. The left external carotid, internal carotid, and occipital arteries were ligated with a 6-0 silk suture, leaving the superior thyroid artery intact. For adenovirus and lentivirus infection studies, adenovirus (Ad-ORF2 or Ad-Scramble) or lentivirus (lenti-shRNA-IRF5 or lenti-shRNA-Scramble) was introduced into the lumen of the left carotid artery and kept inside for 40 min. After infection, the adenovirus or lentivirus was released, and blood flow to the common carotid artery was restored. Mice were fed with a high-fat diet (TD88137, ENVIGO, USA) immediately after surgery and continued for 4 weeks.

### Endothelial AFF3ir-ORF2 overexpression in mice

Endothelial-specific adeno-associated virus (AAV)-mediated CRISPR/Cas9 shuttle plasmid was constructed by Cell & Gene Therapy (Shanghai, China) as previously reported (Wang et al., 2016). ApoE^-/-^ mice received a single tail vein injection of recombinant AAV containing an endothelial-specific human ICAM-2 promoter driving AFF3ir-ORF2 overexpression (AAV- AFF3ir-ORF2) or a control empty vector (AAV-Scramble), with a dose of 1×10^11^ viral genomes in a 200 μL volume of sterile PBS. Subsequently, the mice were fed with a high-fat diet (TD88137, ENVIGO, USA) for 3 months.

### Oil-Red O staining for atherosclerotic plaques in mouse aorta

The ApoE^-/-^, ApoE^-/-^AFF3ir-ORF2^-/-^, and EC-specific AFF3ir-ORF2 overexpression mice were anaesthetized by inhalation of 2% isoflurane and euthanized by cervical dislocation. The aortas were dissected in 1×PBS and opened to expose the atherosclerotic plaques. After fixation in 4% formaldehyde for 1 hat 4 °C, the tissues were rinsed in water for 10 min, followed by 60% isopropanol. The aortas were then stained with Oil-Red O for 30 min with gentle shaking, rinsed again in 60% isopropanol, and subsequently rinsed in water three times. The samples were mounted on wax with the endothelial surface facing upwards. Images were captured using an HP Scanjet G4050. Plaque areas were quantified using NIH ImageJ software by calculating the plaque area relative to the total vascular area.

### Immunofluorescence staining

MEFs slides or frozen sections were fixed in 4% paraformaldehyde for 30 min, then permeabilized in 0.1% Triton X-100 (in PBS) and blocked with 1% bovine serum albumin for 30 min at room temperature. Sections were incubated overnight at 4°C with primary antibodies (1:100). AFF3ir-ORF2 (Cat. No. C0302HL300-4) antibody was from GenScript (Piscataway, NJ, USA). The vWF (Cat. No. ab11713) and VE-Cadherin (Cat. No. ab33168) antibodies were obtained from Abcam (Cambridge, UK). VCAM-1 (Cat. No. sc-13160) antibody was from Santa Cruz Biotechnology (Santa Cruz, CA, USA). Following primary antibody incubation, sections were treated with Alexa Fluor 488- or Alexa Fluor 594-conjugated secondary antibodies (1:200, Thermo Fisher Scientific, Grand Island, NY, USA) at room temperature for 1 hour. Slides were then mounted with DAPI-containing mounting medium. Antibody specificity and target staining authenticity were verified using negative controls. Immunofluorescence micrographs were acquired using a Leica confocal laser scanning microscope. Representative images were randomly selected from each group.

### Histological analysis of atherosclerotic lesions

Harvested carotid arteries and cross-sections of aortic roots were fixed in 4% paraformaldehyde and embedded in optimal cutting temperature compound (OCT). OCT-embedded tissues were sectioned at a thickness of 7 μm. Slides were immersed in 1×PBS for 5 min to remove OCT, and subsequently stained with Oil-Red O, haematoxylin and eosin (HE), and Masson’s trichrome stain to assess lipid accumulation, lesion area, and collagen deposition, respectively (B. Li et al., 2019). Images were acquired using microscopy.

### Quantification of plasma lipid levels

Blood samples were obtained via cardiac puncture, rinsed with heparin, and collected in 1.5 mL Eppendorf tubes. Total plasma cholesterol, triglycerides, LDL cholesterol, and HDL cholesterol levels were measured enzymatically using an automated clinical chemistry analyser kit (Biosino Biotech, Beijing, China).

### Cell culture, transfection, and shear stress experiments

Mouse Embryonic Fibroblasts (MEFs) were obtained and cultured as previously described (Ferreira & Hein, 2023). Cell passages 4 to 7 were used in all experiments. MEFs and HEK293 cells were cultured in the Dulbecco’s Modified Eagle’s Medium (DMEM) supplemented with 10% FBS, penicillin (100 U/mL), and streptomycin (100 μg/mL). Cells were incubated at 37°C in a humidified environment containing 5% CO_2_ and grown to 70-80% confluence before treatment.

Small interfering RNA against IRF5 or IRF8 were synthesized from General Biosystems (Hefei, China). The sequences of siRNAs are shown in the Table S1. The MEFs were passaged to 6- well plated and transfected with 20 nmol/L siRNA per well using the Lipofectamine RNA iMAX Reagent (Invitrogen, Carlsbad, CA, USA).

For flow experiments, confluent monolayers of MEFs were seeded onto glass slides, and a parallel plate flow system was used to launch oscillatory flow (0.5 ± 4 dyn/cm^2^). The flow system was enclosed in a chamber (Frangos, Eskin, McIntire, & Ives, 1985; Fu et al., 2011).

### Adenovirus and lentivirus production and infection

AFF3ir-ORF2 sequences were inserted into the GV138 vector (CMV-MCS-3FLAG) to generate recombinant adenovirus (Ad-AFF3ir-ORF2). The short hairpin RNA (shRNA) sequences targeting mouse IRF5 were 5’-GGGACAACACCATCTTCAAGG-3’, 5’GGTTGCTGCTGGAGATGTTCT-3’, and 5’-GCCTAGAGCAGTTTCTCAATG-3’. The control shRNA was 5’-GCGTGATCTTCACCGACAAGA-3’. These shRNAs were constructed and cloned into pLV-U6-shRNA-CMV-EGFP to generate recombinant lentivirus (lenti-shRNA-IRF5 or lenti-shCtrl). MEFs were infected with adenovirus or lentivirus at a multiplicity of infection (MOI) of 10, with no detectable cellular toxicity observed.

### Western blot analysis

Whole-cell lysates were prepared in a lysis buffer containing a complete protease inhibitor cocktail, PhosSTOP, and PMSF (Roche, Mannheim, Germany). Cytoplasmic and nuclear proteins were extracted from wild-type and AFF3ir-ORF2^-/-^ MEFs using a protein extraction kits (Invent Biotechnologies, SC-003, Beijing, China). Protein were separated by SDS-PAGE and transferred to nitrocellulose membranes (Cat. No. 10600001; GE Healthcare; Chicago, IL, USA). The membranes were incubated with primary antibodies. IRF5 (Cat. No. 96527), IRF8 (Cat. No. 98344), and Flag (Cat. No. 14793) antibodies were from Cell Signaling Technology (Danvers, MA, USA). ICAM-1 (Cat. No. ab222736) antibodies were from Abcam (Cambridge, UK). AFF3ir-ORF2 (Cat. No. C0302HL300-4) and AFF3ir-ORF1 (Cat. No. C0302HL300) antibodies were from GenScript (Piscataway, NJ, USA). GAPDH (Cat. No. 60004-1-Ig) antibody was from Proteintech (Wuhan, China). AFF3 (Cat. No. PA5-68961) antibody was from Thermo Fisher Scientific (Waltham, MA, USA).

After incubation with horseradish peroxidase-conjugated secondary antibodies, the proteins were visualized using enhanced chemiluminescence reagents in a ChemiScope3600 Mini chemiluminescence imaging system (Clinx Science Instruments; Shanghai, China). Protein levels were quantified by measuring integrated density with NIH Image J software (https://imagej.nih.gov/ij/), using GAPDH as a loading control for normalization.

### Co-immunoprecipitation

Whole-cell lysates were prepared by lysing cells in a 1% NP-40 lysis buffer containing 50 mM Tris-HCl, 1% Nonidet-P40, 0.1% SDS, and 150 mM NaCl, supplemented with a complete protease inhibitor cocktail (Cat. No. 04693132001; Roche, Indianapolis, IN, USA), a phosphatase inhibitor (PhosSTOP; Cat. No. 04906845001; Roche), and PMSF (Cat. No. IP0280; Solarbio Life Sciences; Beijing, China). Samples were incubated on ice for 30 minutes, then centrifuged at 12,000 g for 10 minutes, and the supernatant was transferred to a new tube. Protein concentrations were determined using the BCA Protein Assay Kit (Thermo Fisher Scientific, Grand Island, NY, USA).

For immunoprecipitation, 1000 μg of protein was incubated with specific antibodies at 4 °C for 12 hour with constant rotation. Subsequently, 50 μL of 50% Protein A/G PLUS-Agarose beads was added, and the incubation continued for an additional 2 hr. Beads were washed five times with the lysis buffer and collected by centrifugation at 12,000 g for 2 min at 4°C. After the final wash, the supernatant was removed and discarded. Precipitated proteins were eluted by resuspending the beads in 2×SDS-PAGE loading buffer and boiling for 5 min. The eluates from immunoprecipitation were subjected to Western blot analysis.

### ELISA

The concentrations of IL-6 (EM0121) and IL-1β (EM0109) in cell culture supernatant were measured using ELISA kit (FineTest, Wuhan, China). The experiments were conducted according to the protocols provided by the manufacturer.

### Total RNA extraction and real-time quantitative PCR analysis

Total RNA was extracted from cells using an RNA extraction kits (Transgen Biotech, ER501- 01, Beijing, China). Reverse transcription was performed with a reverse transcription kit (Thermo Fisher Scientific, Grand Island, NY, USA). Quantitative PCR was conducted using SYBR Select (Thermo Fisher Scientific) according to the manufacturer’s protocol, with GAPDH serving as the internal control. The primers for quantitative real-time PCR are listed in Supplementary Table 2.

### Luciferase reporter assay

The IRF5-binding motif and the full-length ZNF217 promoter were ligated into pGl3-based plasmids (Genechem, Shanghai, China), as previously described (Qiao et al., 2022). HEK293T cells were seeded into 24-well plates and grown to 70-80% confluency. Cells were transfected with the firefly luciferase reporter plasmid containing the IRF5-responsive ZNF217 promoter along with a β-galactosidase reporter plasmid (Promega, Madison, WI, USA) for 24 hour. Subsequently, cells were then infected with adenovirus (Ad-ORF2 or Ad-Scramble) for an additional 24 hour. Relative luciferase activity was measured using a luciferase assay and normalized to β-galactosidase activity as determined by the β-Galactosidase Enzyme Assay System (Promega, Madison, WI, USA).

### RNA-sequencing (RNA-seq)

RNA-seq was performed as we previously described (Z. Y. Li et al., 2024). Wild-type (WT) and AFF3ir-ORF2^-/-^ MEFs were harvested, and RNA was extracted using the MagicPure Total RNA Kit (TransGen, Beijing, China). Whole transcriptome RNA-seq analysis were conducted by the Beijing Genomics Institute (BGI). Paired-end sequencing in 150-bp length was performed using the DNBSEQ-G400 platform. Raw data was filtrated using SOAPnuke (v1.5.6). Differential gene expression analysis, with thresholds set at *P* < 0.05 and fold change ≥1.5, was performed via the BGI website (http://omiscribe.bgi.com). Pathway enrichment analysis was carried out using DAVID tools.

### Statistical analysis

Statistics analyses were performed using GraphPad Prism 8.0. No sample outliers were excluded. At least six independent experiments were performed for all biochemical experiments and the representative images were shown. Unpaired Student’s *t* test (2-tailed), 1-way ANOVA or 2-way ANOVA with Bonferroni multiple comparison post hoc test were used for analyses, as appropriate. Sample size, statistical method, and statistical significance are specified in Figures and Figure Legends. Levels of probabilities less than 0.05 were regarded as significant.

## Data Availability Statement

The data that support the findings of this study are available from the corresponding author upon reasonable request. The RNA-sequencing data has been submitted to GEO datasets (GSE286206). Source data are provided with this paper.

## Acknowledgments

This work was supported by National Natural Science Foundation of China Grants (82330012, 82127808, 82422006, 32471166, and 82270516), British Heart Foundation (FS-15/74/31669), Natural Science Foundation of Tianjin (24JCJQJC00060 and 21JCYBJC01590), and the Key Research Project in Traditional Chinese Medicine of Tianjin (2023013).

