## supplemental figure 1-7 and supplemental table 1-2 for "Disruption of the Novel Nested Gene Aff3ir Mediates Disturbed Flow-Induced Atherosclerosis in Mice"

**Running title:** AFF3ir-ORF2 exerts anti-atherosclerosis roles

[#]These authors contributed equally to this study.

***Correspondence:**

Jinlong He, Ph.D.

Department of Physiology and Pathophysiology, Tianjin Medical University; Qixiangtai Rd 22nd, Tianjin, 300070, China.

or

Yi Zhu, M.D., Ph.D.

Department of Physiology and Pathophysiology, Tianjin Medical University; Qixiangtai Rd 22nd, Tianjin, 300070, China.

or

Lingfang Zeng, Ph.D.

School of Cardiovascular and Metabolic Medicine and Sciences, Faculty of Life Science and Medicine, King's College London, London SE5 9NU, United Kingdom.

**Keywords:** Shear stress, Atherosclerosis, Nested gene, IRF5

### Supplementary Figures and Figure legends

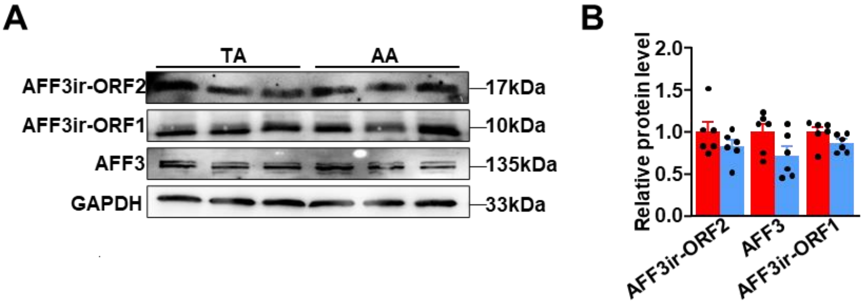


**Figure S1. A-B,** Media and adventitia of aortas were isolated from the C57BL/6 mice. Western blot analysis of the expression of the indicated proteins in the media and adventitia of TA (thoracic aorta) and AA (aortic arch). Protein levels were normalized to GAPDH, and the relative expression values were compared to the TA group. Data are presented as mean ± SEM (n=6 mice per group). **P*< 0.05, unpaired two-tailed *t*-test.

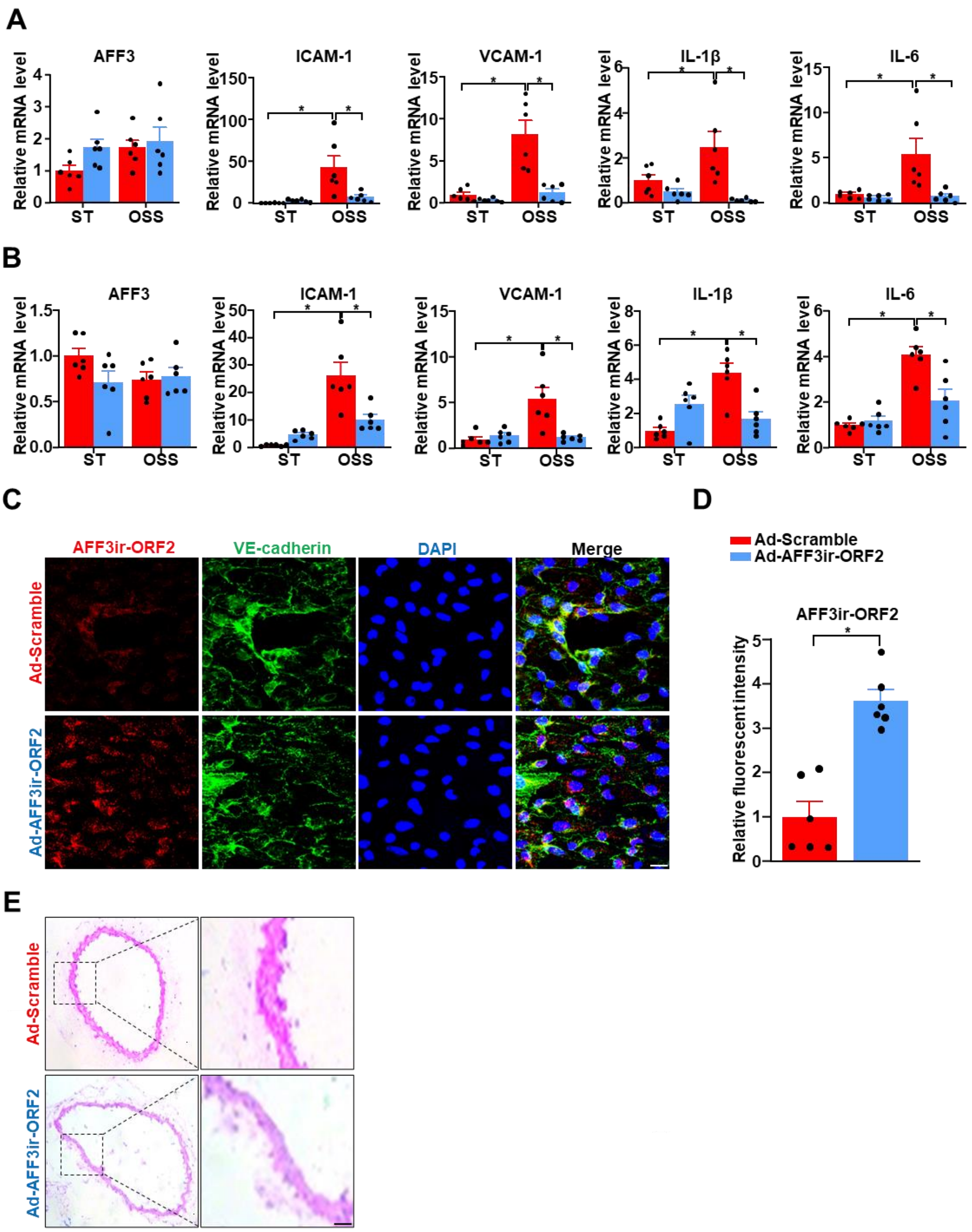


**Figure S2. A,** Mouse embryonic fibroblasts (MEFs) isolated from C57BL/6 mice were infected with indicated adenoviruses (Ad-Scramble or Ad-Aff3ir-ORF2) for 48 h and then exposed to static (ST) or oscillatory shear stress (OSS, 0.5 ± 4 dyn/cm$^2$, 1 Hz) for another 6 h. RT-PCR analysis of mRNA levels of AFF3, VCAM-1, ICAM-1, interleukin-6 (IL-6), and interleukin-1 beta (IL-1β). The relative expression values were compared to MEFs infected with Ad-Scramble and treated with ST. Data are presented as mean ± SEM (n = 6 independent experiments). *P< 0.05, two-way ANOVA with Tukey post-test. **B,** Human umbilical vein endothelial cells (HUVECs) were infected with Ad-Scramble or Ad-Aff3ir-ORF2 for 48 h and then exposed to ST or OSS for another 6 h. RT-PCR analysis of mRNA levels of AFF3, VCAM-1, ICAM-1, IL-6, and IL-1β. The relative expression values were compared to HUVECs infected with Ad-Scramble and treated with ST. Data are presented as mean ± SEM (n = 6 independent experiments). **P*<0.05, two-way ANOVA with Tukey post-test. **C-D,** Enface immunofluorescence staining of AFF3ir-ORF2, VE-cadherin, and DAPI and quantification of AFF3ir-ORF2 expression. Scale bar, 20 μm. The immunofluorescence intensity of AFF3ir-ORF2 was normalized to DAPI and the relative expression values were compared to that of the Ad-Scramble group. Data are presented as mean ± SEM (n=6 mice per group). **P*< 0.05, unpaired two-tailed *t*-test. **E**, Eight-week-old male $ApoE^{-/-}$ mice were subjected to partial ligation of the carotid artery along with 10 μL of adenovirus suspension at $1 \times 10^8$ transducing units (TU)/mL was instilled into the left carotid arteries. The mice were then fed high-fat diet for 4 weeks. Right carotid arteries were sectioned for hematoxylin and eosin staining.

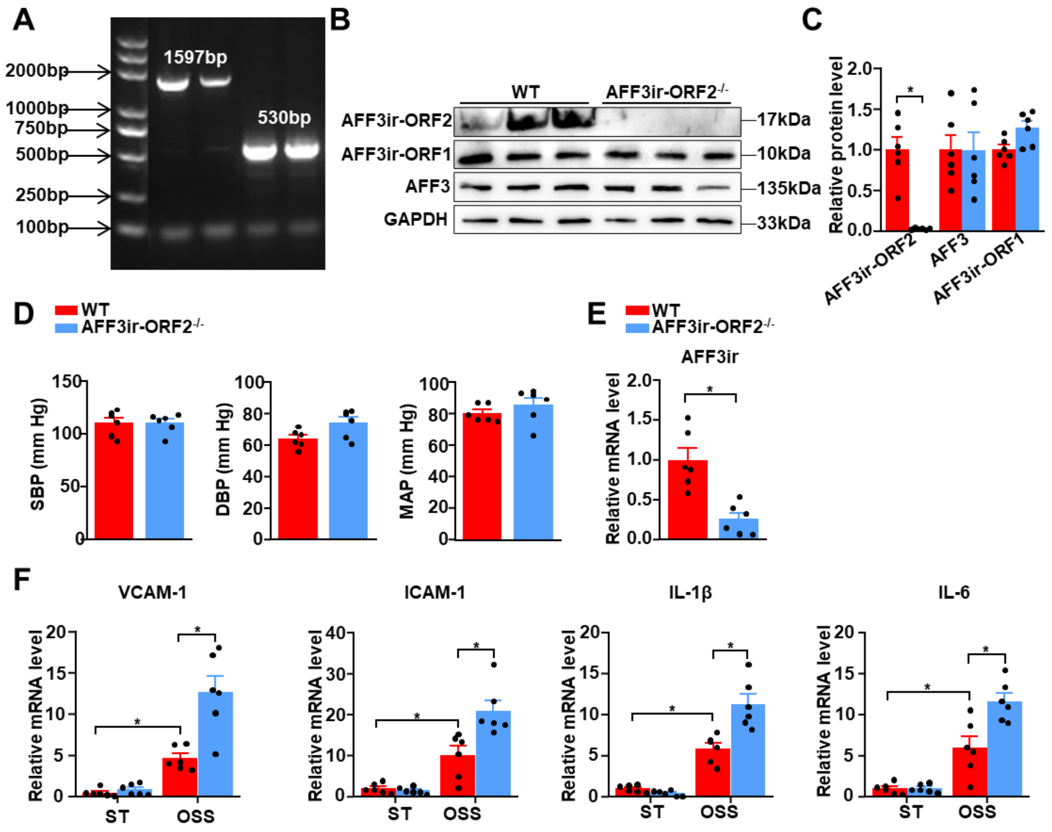


**Figure S3. A,** Genetic typing of the AFF3ir-ORF2 mice by the PCR-RFLP assay. Lane 1: Marker; Lanes 2-3: Wild type (1597 bp); Lanes 4-5: AFF3ir-ORF2$^{-/-}$ (530 bp). **B-C,** Western blot analysis of the expression of the indicated proteins in the intima of WT and AFF3ir-ORF2$^{-/-}$ mice. Protein levels were normalized to GAPDH, and relative expression values were compared to the WT group. Data are presented as mean ± SEM (n=6 mice per group). **P*<0.05, unpaired two-tailed *t*-test. **D,** Quantification of systolic blood pressure (SBP), diastolic blood pressure (DBP), and mean artery pressure (MAP) in eight-week-old male wild-type (WT) and AFF3ir-ORF2$^{-/-}$ mice. Data are presented as mean ± SEM (n=6 mice per group). **P*< 0.05, unpaired two-tailed *t*-test. **E,** Mouse embryonic fibroblasts (MEFs) isolated from WT and AFF3ir-ORF2$^{-/-}$ mice, and RT-PCR analysis of the mRNA levels of *AFF3ir-ORF2*. Data are presented as mean ± SEM (n=6 mice per group). **P*< 0.05, unpaired two-tailed *t*-test. **F,** MEFs isolated from WT and AFF3ir-ORF2$^{-/-}$ mice were exposed to static (ST) or oscillatory shear stress (OSS, 0.5 ± 4 dyn/cm$^2$, 1 Hz) for 6 h. RT-PCR analysis of the mRNA levels of *AFF3ir-ORF2*, *ICAM-1*, and *VCAM-1*, *IL-6*, and *IL-1β*. The relative expression values were compared to that of WT MEFs treated with ST. Data are presented as mean ± SEM (n = 6 independent experiments). **P*<0.05, two-way ANOVA with Tukey post-test.

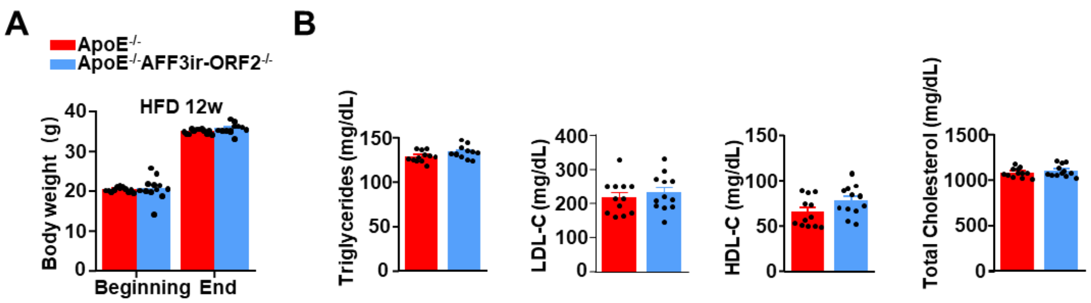


**Figure S4.** Eight-week-old male ApoE$^{-/-}$ and ApoE$^{-/-}$ORF2$^{-/-}$ mice were fed a high-fat diet (HFD) for 12 weeks. **A,** Quantification of body weight at the beginning and end of the experiment. Data are presented as mean ± SEM (n=12 mice per group). **P*< 0.05, two-way ANOVA with Tukey post-test. **B,** Quantification of plasma levels of triglycerides, total cholesterol, low-density lipoprotein cholesterol (LDL-C), and high-density lipoprotein cholesterol (HDL-C). Data are presented as mean ± SEM (n=12 mice per group). **P*< 0.05, unpaired two-tailed *t*-test.

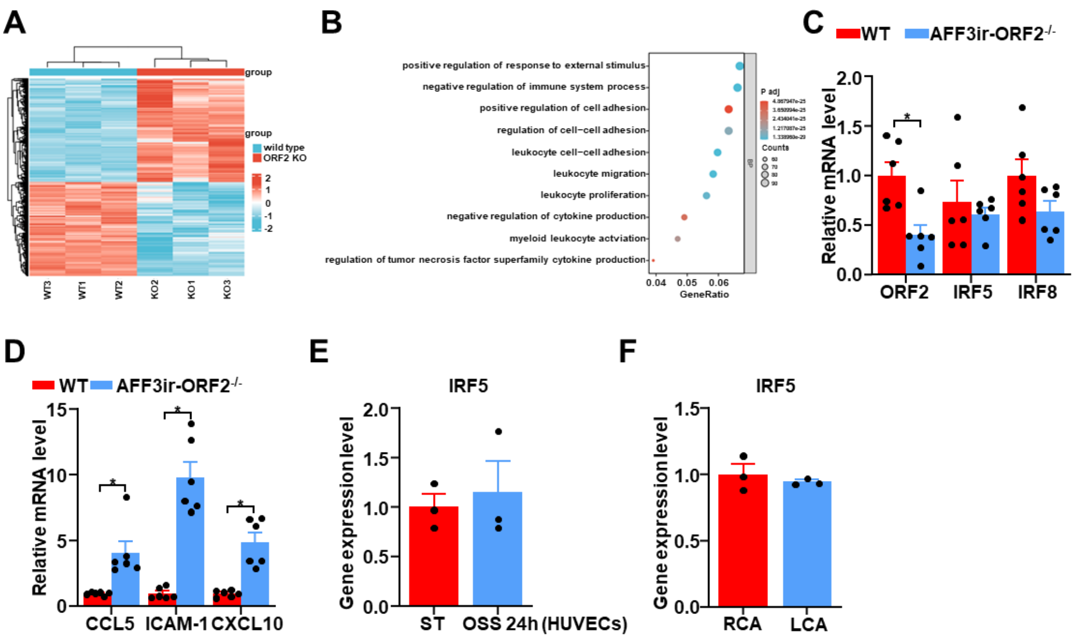


**Figure S5. A,** Heat map showing profiles of all the differentially expressed genes from the RNA-seq analysis of wild-type (WT) and AFF3ir-ORF2$^{-/-}$ mouse embryonic fibroblasts (MEFs) (n=3). **B,** All the differentially expressed genes from the RNA-seq data were mapped onto the atherosclerosis-related gene dataset. The overlapping genes were then subjected to Gene Ontology enrichment pathway analysis. **C**, RT-PCR analysis of the mRNA levels of ORF2, IRF5, and IRF8 in WT and AFF3ir-ORF2$^{-/-}$ MEFs. The relative expression values were normalized to WT MEFs. Data are mean ± SEM (n = 6 independent experiments). *$P$< 0.05, unpaired two-tailed *t*-test. **D,** RT-PCR analysis of the mRNA levels of CCL5, ICAM-1 and CXCL10 in WT and AFF3ir-ORF2$^{-/-}$ MEFs. The relative expression values were compared to WT MEFs. Data are mean ± SEM (n = 6 independent experiments). *$P$< 0.05, unpaired two-tailed *t*-test. **E**, IRF5 mRNA levels in HUVECs under static (ST) and oscillatory shear stress (OSS, 0.5 ± 4 dyn/cm$^2$, 1 Hz) condition for 24 hours were shown. The mRNA expression levels were sourced from the GEO database (GSE276195). Data are mean ± SEM (n = 3). **F**, Partial carotid ligation was performed on the left carotid artery (LCA) of the mouse while the right carotid artery (RCA) was left untouched. IRF5 mRNA levels in LCAs and RCAs were shown. The mRNA expression levels were sourced from the GEO database (GSE222583). Data are mean ± SEM (n = 3).

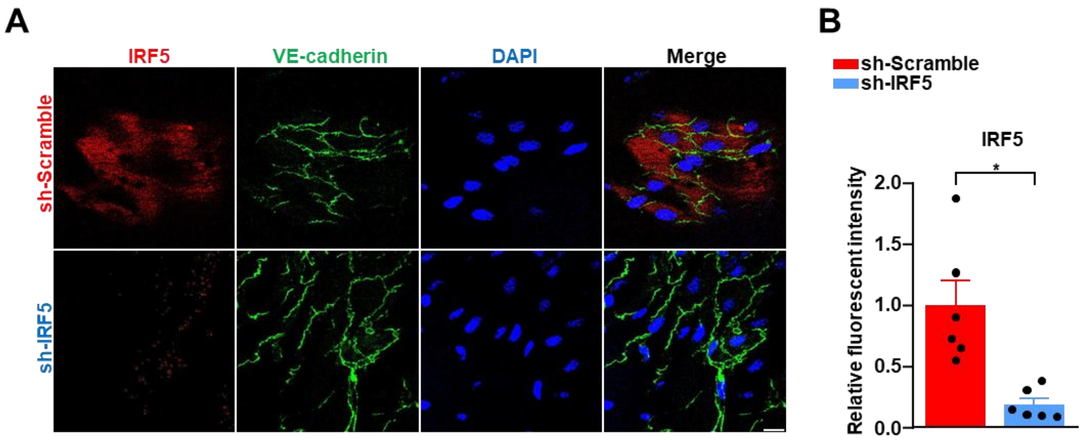


**Figure S6. A-B,** Enface immunofluorescence staining of IRF5, VE-cadherin, and DAPI and quantification of IRF5 expression. Scale bar, 20 μm. The immunofluorescence intensity of IRF5 was normalized to DAPI, and the relative expression values were compared to that of sh-Scramble group. Data are presented as mean ± SEM (n=6 mice per group). **P*< 0.05, unpaired two-tailed *t*-test.

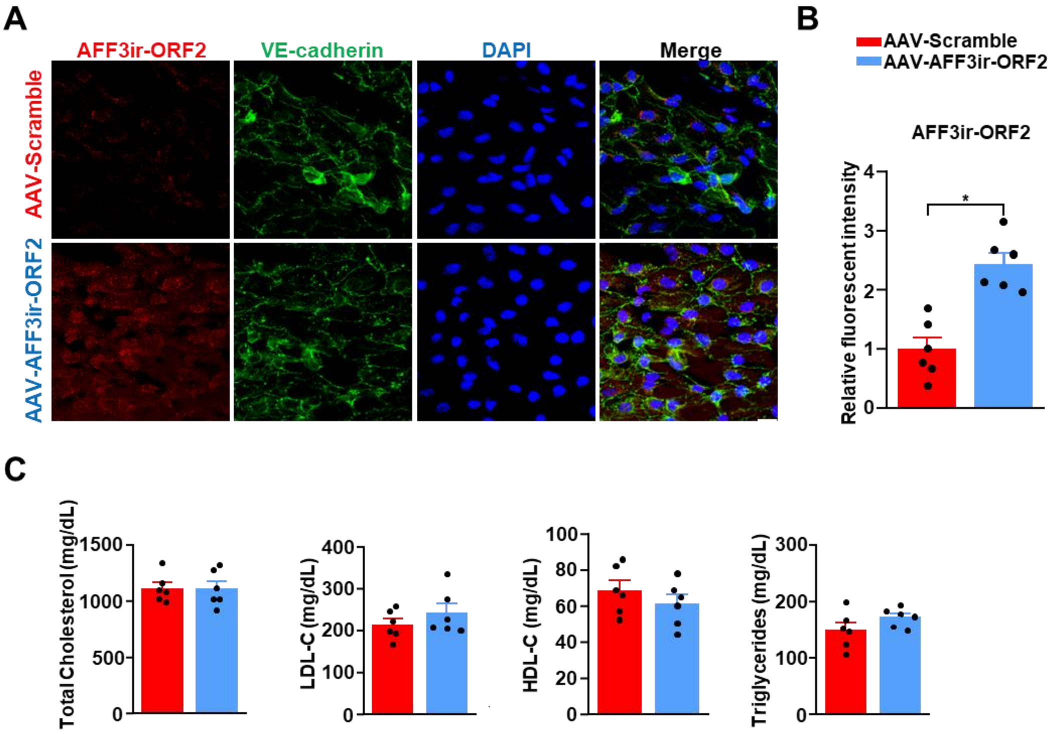


**Figure S7. A-B,** Enface immunofluorescence staining of AFF3ir-ORF2, VE-cadherin, and DAPI and quantification of AFF3ir-ORF2 expression. Scale bar, 20 µm. The immunofluorescence intensity of AFF3ir-ORF2 was normalized to DAPI, and the relative expression values were compared to that of the AAV-Scramble group. Data are presented as mean ± SEM (n=6 mice per group). *$P$< 0.05, unpaired two-tailed $t$-test. **C,** Quantification of plasma levels of triglycerides, total cholesterol, low-density lipoprotein cholesterol (LDL-C), and high-density lipoprotein cholesterol (HDL-C). Data are presented as mean ± SEM (n=6 mice per group). *$P$< 0.05, unpaired two-tailed $t$-test.

**Table S1. The sequences of siRNAs.**

| siRNA | Stand | Sequence |
|---|---|---|
| siIRF5-1 | Forward | GGAAAAGAAACUCUUCUAUTT |
| | Reverse | AUAGAAGAGUUUCUUUUCCTT |
| siIRF5-2 | Forward | CACCUUUUGAGAUCUUCUUTT |
| | Reverse | AAGAAGAUCUCAAAAGGUGTT |
| siIRF8-1 | Forward | GUUUAAAGAGGGAGACAAATT |
| | Reverse | UUUGUCUCCCUCUUUAAACTT |
| siIRF8-2 | Forward | CCAACAAGCUGGAGCGGGATT |
| | Reverse | UCCCGCUCCAGCUUGUUGGTT |
| siIRF8-3 | Forward | GAGCGAAGUUCCUGAGAUGTT |
| | Reverse | CAUCUCAGGAACUUCGCUCTT |

**Table S2. Primers for qRT-PCR.**

| Gene | Stand | Sequence |
|---|---|---|
| GAPDH | Forward<br>Reverse | AGGTCGGTGTGAACGGATTTG<br>TGTAGACCATGTAGTTGAGGTCA |
| IRF5 | Forward<br>Reverse | AGAGACAGGGAAGTACACTGAAG<br>TGGAAGTCACGGCTTTTGTTAAG |
| IRF8 | Forward<br>Reverse | CGGGGCTGATCTGGGAAAAT<br>CACAGCGTAACCTCGTCTTC |
| AFF3 | Forward<br>Reverse | TCTGTGGGCTCCATCAAC<br>GACCTCATCGCTGTCCTTT |
| AFF3ir | Forward<br>Reverse | GAAACTGAACACGGGAGG<br>GAACCTGATGCCTGGAAC |
| VCAM-1 | Forward<br>Reverse | AGTTGGGGATTCGGTTGTTCT<br>CCCCTCATTCCTTACCACCC |
| ICAM-1 | Forward<br>Reverse | GTGATGCTCAGGTATCCATCCA<br>CACAGTTCTCAAAGCACAGCG |
| IL-6 | Forward<br>Reverse | TAGTCCTTCCTACCCCAATTTCC<br>TTGGTCCTTAGCCACTCCTTC |
| IL-1β | Forward<br>Reverse | GCAACTGTTCCTGAACTCAACT<br>ATCTTTTGGGGTCCGTCAACT |
